# Temporal and sequential transcriptional dynamics define lineage *shifts in corticogenesis*

**DOI:** 10.1101/2022.02.10.479992

**Authors:** Tanzila Mukhtar, Jeremie Breda, Marcelo Boareto, Pascal Grobecker, Zahra Karimaddini, Alice Grison, Katja Eschbach, Ramakrishnan Chandrasekhar, Swen Vermeul, Michal Okoniewski, Mikhail Pachkov, Suzana Atanasoski, Christian Beisel, Dagmar Iber, Erik van Nimwegen, Verdon Taylor

## Abstract

The cerebral cortex contains billions of neurons, and their disorganization or misspecification leads to neurodevelopmental disorders. Understanding how the plethora of projection neuron subtypes are generated by cortical neural stem cells (NSCs) is a major challenge. Here, we focused on elucidating the transcriptional landscape of murine embryonic NSCs, basal progenitors (BPs) and newborn neurons (NBNs) throughout cortical development. We uncover dynamic shifts in transcriptional space over time, and heterogeneity within each progenitor population. We identified signature hallmarks of NSC, BP and NBN clusters, and predict active transcriptional nodes and networks that contribute to neural fate specification. We find that the expression of receptors, ligands and downstream pathway components is highly dynamic over time and throughout the lineage implying differential responsiveness to signals. Thus, we provide an expansive compendium of gene expression during cortical development that will be an invaluable resource for studying neural developmental processes and neurodevelopmental disorders.

## Introduction

The cerebral cortex of vertebrates is an isocortex, composed of six layers of morphologically and functionally distinct neurons. During development, cortical NSCs pass through consecutive stages of mitotic expansion, deep- to upper-layer neurogenesis and then gliogenesis. Most neurons are generated from NSCs through a transient progenitor population, the BPs. Maintenance of progenitor potential and control of cortical fate commitment are regulated through the integration of dynamic signaling pathways organized in space and time, which induces an elaborate interplay between downstream transcriptional networks. Although the molecular nature of mature neurons within the six cortical layers has been described, their corresponding progenitors have not been clearly characterized.

Different hypotheses have been proposed to explain the heterogeneity in the cortical precursor cells in terms of temporal expansion and differentiation potential (Hevner et al., 2003; Lodato and Arlotta, 2015; Molyneaux et al., 2007; Woodworth et al., 2012). One hypothesis states that NSCs switch their fate temporally in coherence with the time points of neurogenesis and thus generate neurons of successive layers of the cortex followed by glial cells (Guo et al., 2013). An alternate hypothesis proposes that NSCs are a multipotent cell pool, wherein each cell would be guided by intrinsic and extrinsic signals to generate a specific selection of neuronal subtypes or glial cells and these different progenitors are recruited in a sequential manner (Franco et al., 2012). Whether one or both hypotheses are correct remains a major debate.

As RNA sequencing (RNA-Seq) technology increased over recent years, so has our acceptance of an increasing repertoire of cell types present during cortical development. Particularly single cell sequencing techniques have allowed an ever more detailed transcriptomic analysis of cortical precursor cells (Arber et al., 2015; Arlotta et al., 2005; Chuang et al., 2015; Desai and McConnell, 2000; Di Bella et al., 2021; Ecker et al., 2017; Fode et al., 2000; Gotz and Huttner, 2005; Greig et al., 2013; Han and Sestan, 2013; Haubensak et al., 2004; Hevner *et al*., 2003; Johnson and Walsh, 2017; Liu et al., 2016; Lodato and Arlotta, 2015; Lui et al., 2011; Molyneaux *et al*., 2007; Mukhtar and Taylor, 2018; Nowakowski et al., 2017; Paridaen and Huttner, 2014; Pollen et al., 2015; Rosenberg et al., 2018; Stancik et al., 2010; Telley et al., 2019a; Telley et al., 2016). Frequently, cells are isolated based on positional information or temporal labelling and this is used to delineate cell-type and predict potential (Di Bella *et al*., 2021; Telley et al., 2019b). Although these approaches have been very successful in providing a framework, our understanding of transcriptional programs during brain development and cortical patterning is not complete and some critical points remain. One major challenge is the extreme complexity of the system and the differences in technical approaches undertaken. As RNA-Seq takes a snapshot in time of gene expression in a population or of single cells, it is challenging to predict the past and future gene expression profile of a cell population. Elegant labelling procedures have provided some insight into cell diversity in the NSC pool and allowed analysis of specific gene function (Telley *et al*., 2019a; Telley *et al*., 2016). However, it remains unclear how gene expression within the defined populations of NSCs and progenitors in the developing mammalian cortex *in vivo* change over time and through the lineage as the fate decisions are being made.

In order to compare like-with-like and circumvent some of the challenges of random cell selection, we took advantage of the knowledge about murine cortical development and transgenic mice that allow isolation of defined progenitor populations at each day between embryonic day 10.5 (E10.5) and birth (Hebert and Fishell, 2008). We performed bulk and single cell RNA-Seq to generate gene expression profiles of NSCs, BPs and NBNs from the dorsal cortex, spanning the critical periods of NSC expansion (E10.5-11.5), neurogenesis (E12.5-16.5) and gliogenesis (E17.5-PN1). From these data catalogues, we elucidated the transcriptional landscapes of NSCs, BPs and neuronal subtypes and systematically followed robust temporal dynamics in their gene expression through cortical development. We determined an amazing dynamic heterogeneity within these progenitor populations at the single-cell level, identifying clusters of NSCs, BPs and NBNs and providing gene signatures for each cluster. We evaluated the changes in signaling pathway component expression during cortical development and identified receptors, ligands and downstream signaling pathways that potentially play critical roles in cortical development. Finally, we found that the transcriptional programs that define specific cortical neuron type, are active in NSCs prior to the birth of the neurons. Our work provides a versatile and comprehensive resource that will be useful to address gene expression but also novel aspects of NSCs fate choice and neuronal cell subtype generation.

## Results

### Transcriptional analyses validate the selection and sorting procedure

Canonical Notch signaling in the developing cortex suppresses NSC differentiation by repressing expression of proneurogenic transcription factors while promoting proliferation and survival (Dang et al., 2006; Gaiano and Fishell, 2002; Imayoshi et al., 2010; Kageyama et al., 2009; Mason et al., 2006). *Hes5* is a transcriptional target of Notch signaling and labels NSCs at all stages of development and in the adult (Bansod et al., 2017; Basak et al., 2012; Lugert et al., 2010; Lugert et al., 2012). Conversely, *Eomes* (*Tbr2*) is expressed by BPs and committed neural progenitors (Arnold et al., 2008; Sessa et al., 2017). *Hes5::GFP* labels NSCs in the ventricular zone and *Tbr2::GFP* BPs and NBNs in the subventricular zone and developing cortical plate (Figures S1A,B) (Arnold et al., 2009; Basak and Taylor, 2007).

To address changes in gene expression within the NSC, BPs and early neurons of the cortical neural lineages, cells were sorted from individual *Hes5::GFP* and *Tbr2::GFP* embryos at each day of development between embryonic day 10.5 (E10.5) and birth (PN) and RNA-Seq performed on the samples from each embryo separately (Figures 1A and S1A). E10.5- PN1 covered the embryonic stages of cortical development from NSC expansion (E10.5- E11.5), through neurogenesis (E12.5-E16.5) to gliogenesis (E17.5-PN).

**Figure 1:**
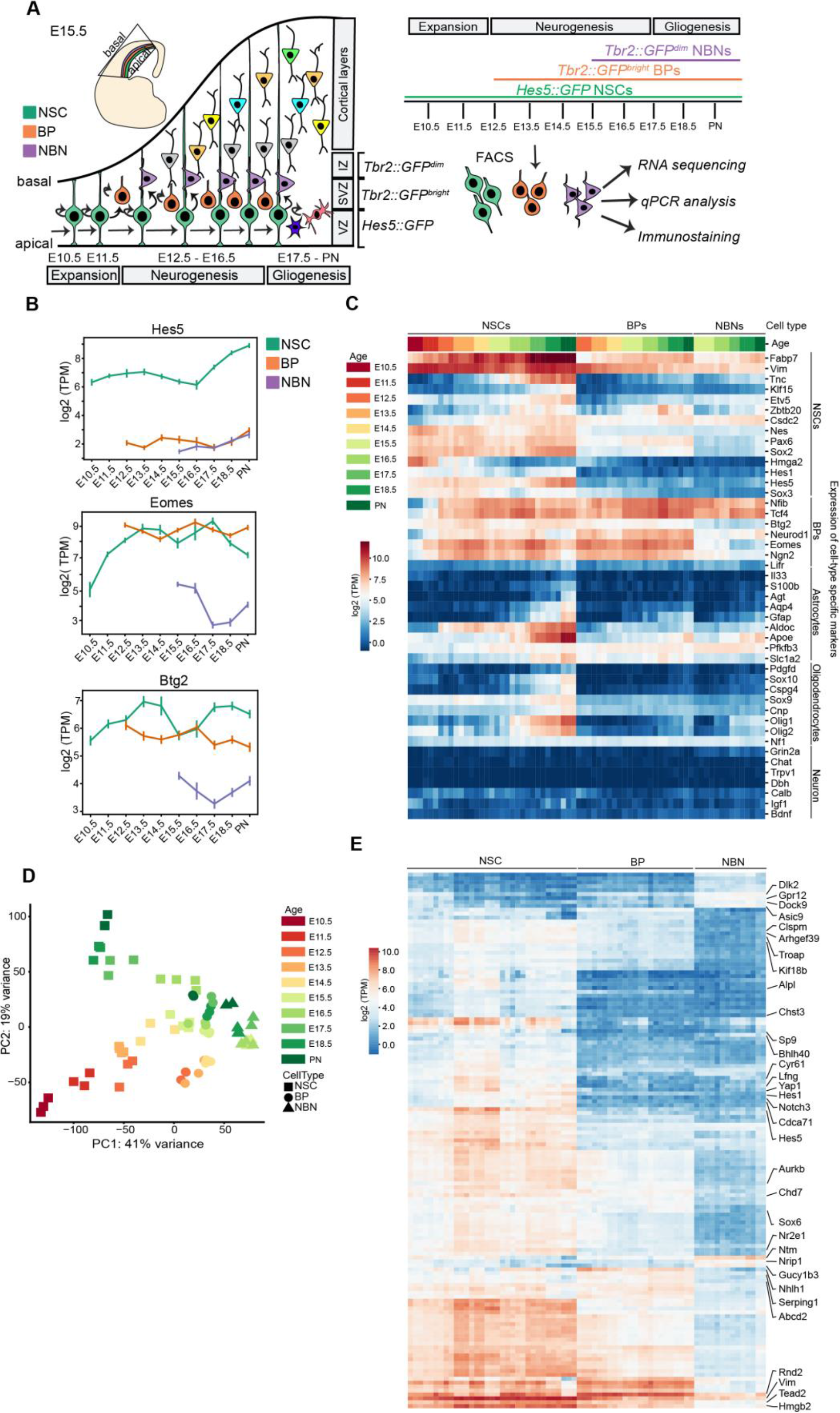
Overview and validation of the transcriptional analyses. (A) Overview of the biological system with experimental paradigm, illustrating NSCs, BPs and NBNs were isolated at each day during development from E10.5 to PN. (B) Notch signaling effector *Hes5* is expressed high in NSCs while *Eomes (Tbr2)* and *Btg2* are expressed high in both NSCs and BPs at the mRNA level. (C) Heatmap validating the known cell-type specific marker gene expression from RNA sequencing data. (D) Principal Component Analysis (PCA) for all samples of NSCs, BPs and NBNs throughout development, covering maximum variance. (E) Heatmap illustrating the novel marker genes identified from the RNA sequencing data, as signature genes for NSCs, BPs and NBNs. NSCs- Neural stem cells, BPs-Basal progenitors, NBNs-Newborn neurons, E-Embryonic day, PN-Post natal, VZ-Ventricular zone, SVZ- Subventricular zone, IZ-Intermediate Zone. Expression values on the heatmaps are log2 (transcripts per million).

We gated on *Hes5::GFP^high^* cells at all time points excluding low and negative cells (Figures S1C). Immunostaining of acutely sorted *Hes5::GFP^+^* cells showed expression of the NSC associated proteins Sox2 and Pax6 but not Tbr2 (Figures S1D). *Tbr2::GFP^+^* cells were first detectable by FACS at E12.5 corresponding to the prominent appearance of BPs in the developing dorsal cortex (Arnold *et al*., 2009). From E15.5 on, the *Tbr2::GFP^+^* population was divided into GFP^high^, GFP^low^ and GFP^-^ populations (Figures S1C). We separated *Tbr2::GFP^low^* and *Tbr2::GFP^high^* cells and analyzed these populations separately. Sorted *Tbr2::GFP^high^* cells expressed low levels of Sox2 and Pax6 and high levels of Tbr2 denoting them as BPs (Figures S1E). Sorted *Tbr2::GFP^low^* cells did not express Sox2 and Pax6 and had lower levels of Tbr2 than the BPs (Figures S1F). We reasoned that the *Tbr2::GFP^low^* cells were immature NBNs labelled by low levels of *Tbr2*and perduring GFP.

As proof of concept, *Hes5* RNA levels were high in the NSCs populations at all developmental time points and low in the *Tbr2::GFP^+^* samples (Figure 1B). As expected, the transcripts of the BP markers *Tbr2* and *Btg2* were highly expressed by *Tbr2::GFP^high^* cells at all stages from E12.5-PN1 consistent with being dorsal cortical BPs and at lower levels by *Tbr2::GFP^low^* NBNs (Figure 1B). *Tbr2* and *Btg2* transcripts were detected in the NSC samples without detectable protein, confirming previous observations (Figures 1B and S1B, D-F) (Mukhtar et al., 2020; Pollen *et al*., 2015). These points demonstrate the challenges of allocation of cell-type based on transcriptional activity of a few “marker” genes.

Interrogation of the RNA-Seq data revealed that NSCs, BPs and NBN transcriptomes were remarkably similar, and few genes were differentially expressed between NSCs and BPs (Figures 1C). We analyzed the dynamics in expression of known NSC, BP and NBN markers between E10.5 and PN1 (Figures 1C, S1G and S1H). NSC markers were highly expressed throughout cortical development with characteristic temporal dynamics in the NSC populations (Gotz and Huttner, 2005; Molyneaux *et al*., 2007; Mukhtar and Taylor, 2018; Ohtsuka et al., 2011; Pollen *et al*., 2015).

Known BP markers, including *Nfib, Ngn2, Tcf4,* and *Neurod1,* were highly expressed throughout cortical development by BPs. Astrocytic markers including *S100b, ApoE, Gfap,* and *Aldoc* were expressed highly by NSCs isolated late in development corresponding to the onset of gliogenesis indicating that the glial transcriptional program had already been initiated (Liddelow and Barres, 2015; Molofsky et al., 2012; Zhang and Barres, 2010). Similarly, key markers for oligodendrocytes including *Pdgfd, Sox10, Cspg4,* and *Sox9* were expressed higher by late stage NSCs corresponding to the last wave of oligodendrogenesis originating in the ventricular zone of the dorsal cortex (Ono et al., 2008; Takebayashi and Ikenaka, 2015; Zhang and Barres, 2010). The mature neuronal markers *Grin2a, Chat, Bdnf,* and *Igf1* were expressed at very low levels by the *Hes5::GFP* and *Tbr2::GFP* sorted cells indicating that the selection process isolated progenitors and excluded mature neurons (Figures 1C and S1G) (Sarnat, 2013).

Unbiased computational analyses revealed extensive transcriptional dynamics within the different cell types. Principal component analysis (PCA) capturing 60% of the total variance (PC1 and PC2) separated the samples based on cell type (NSC, BP and NBN) and developmental stage. Projection of the BPs and NBNs onto the first two PCs indicated they are transcriptionally closer to NSCs in the neurogenic phase of cortical development than those in the expansion and gliogenic phases (Figures 1D and S1I). The BPs were positioned between the NSCs and the NBNs in the neurogenic phase consistent with being a transient neuronal precursor population (Figures 1D and S1I). NSCs displayed maximum variations in gene expression across time on the first two PCs, with a fluid separation from the expansion to neurogenesis and gliogenesis phases. Subsequently, we performed pairwise differentially expressed gene analyses (DEG) between the different cell types (Figures 1E and S1J-O). To exclude substantial contamination of the *Hes5::GFP^high^* sorted cells with BPs and NBNs, we identified transcripts that were highly expressed by BPs and NBNs but not by NSCs (Figure S1J-O). Thus, although the transcriptomes of NSCs and BPs are remarkably similar, the sorting procedures were effective in enriching stem from progenitor populations throughout cortical development. We identified novel markers for NSCs, BPs and NBNs using two independent methods - DEGs and Z-score log2 (TPM) expression values (Table S1). Genes including *Sp9, Cyr61, Yap1, Hes1, Lfng,* and *Notch3* are highly expressed by NSCs. Identification of these signature genes using an unbiased approach is and independent validation of the approach as the function of some have been studied in NSCs. The Hippo co-activator *Yap1*, Notch signaling components *Hes1*, *Lfng* are involved in NSC proliferation and maintenance (Bray, 2006; Pourquie, 2003; Takebayashi and Ikenaka, 2015). *Gucy1b3, Nhlh1, and Serping1* were highly expressed by BPs and novel markers of the cell-type in the lineage but not much is known about their function (Lipkowitz et al., 1992). *Ntm, Nrip1* are expressed higher in NBNs than in BPs or NSCs also providing novel markers (Gil et al., 1998). Interestingly, DEG analyses revealed that the majority of the highly expressed genes in NSCs are downregulated by BPs and reduced further by NBNs.

### Temporal dynamics in transcriptional landscapes of NSCs, BPs and NBNs based on gene expression

Our analyses showed that NSCs displayed maximum variance over time and therefore contribute heavily to the first two PCs. To understand the transcriptional dynamics in the NSCs, we performed PCA focusing only on the NSCs. The first two PCs covered almost 70% of the total variance and exposed a dynamic transcriptional path among the phases of expansion, neurogenesis and gliogenesis (Figures 2A and S2A). Although the NSCs were isolated using the same characteristic, *Hes5::GFP^high^* expression, we observed striking, stage-related dynamic movement through transcriptional space. PCA indicated that NSCs could follow a continuous path from expansion through neurogenesis to gliogenesis, consistent with the common origin model of sequential cell specification over time. Surprisingly, *Hbb-bh1, Hba-x* and *Hbb-y* expression distinguished NSCs in the expansion phase (PC1 negative axis) from those in the neurogenic and gliogenic phases (Figure 2B). Although hemoglobin subunits are predominantly associated with erythrocytes and oxygen transport from the lungs, hemoglobin subunits are also expressed in the brain and by neurons (Brown et al., 2016). Hemoglobin subunits are found in the mitochondria in neurons and may assist oxygen transport across mitochondria membranes. As NSCs are mitotically highly active and require extensive energy for cell division, it is possible that hemoglobin supports the energy requirements during the expansion phase.

**Figure 2:**
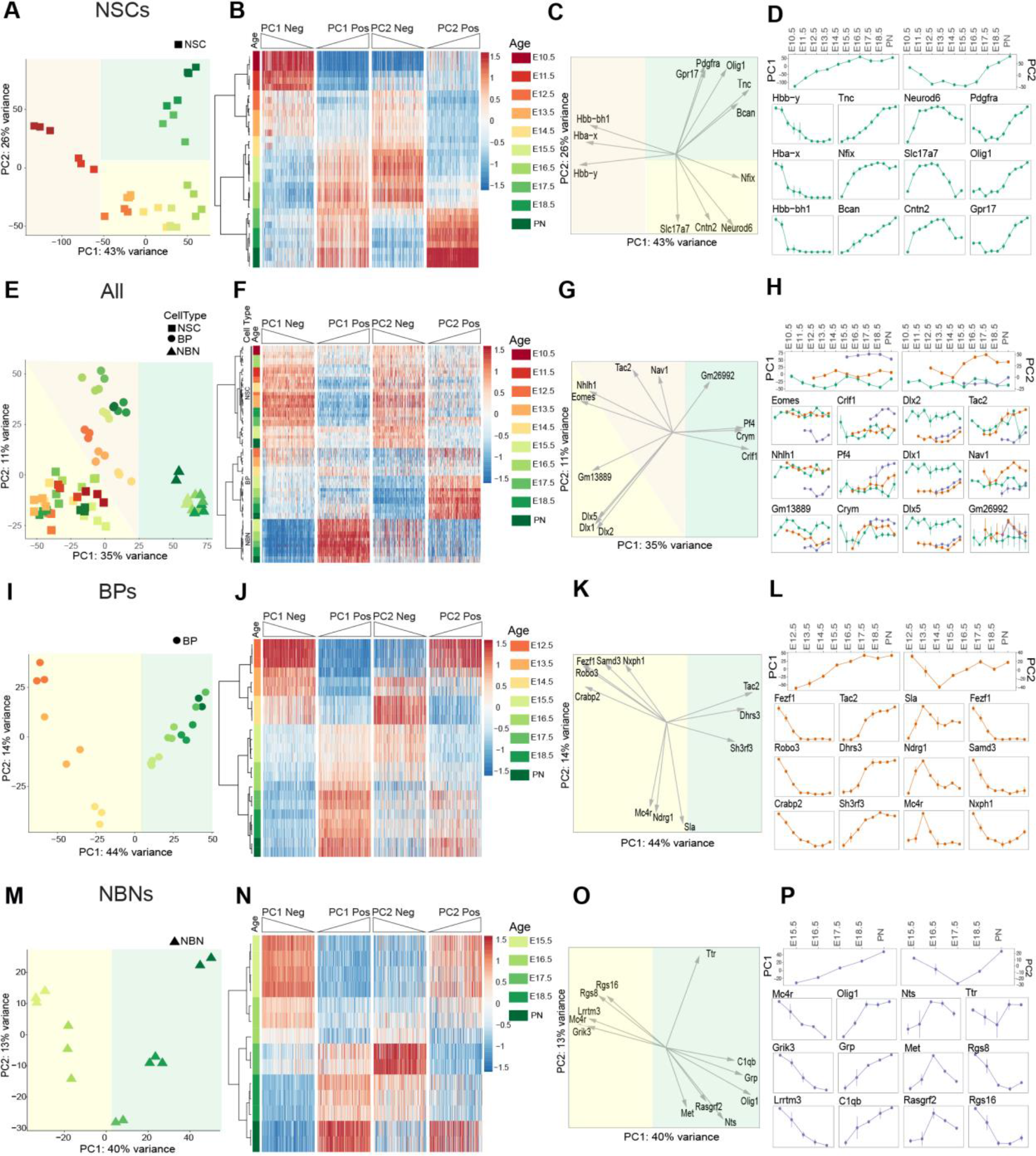
Dynamics of transcriptional profile changes in different populations over time. (A) PCA of NSCs from E10.5 to PN showing their transcriptional dynamics. (B) Heatmap of genes that have the highest contribution to the PC1 and PC2 for NSCs, sorted by their weights (250 genes from each side). (C) PCA plots with projected genes shown as vectors illustrating their contribution. (D) Illustrating the trends (top), based on the position of samples along PC1 and PC2, and the gene expression profiles of top three genes from each side of PC1 and PC2 for NSCs (bottom). (E) PCA of all samples removing the first two principal components of NSCs from E10.5 to PN showing their transcriptional dynamics. (F) Heatmap of genes that have the highest contribution to the PC1 and PC2, sorted by their weights (250 genes from each side). (G) PCA plots with projected genes shown as vectors illustrating their contribution. (H) Illustrating the trends (top), based on the position of samples along PC1 and PC2, and the gene expression profiles of top three genes from each side of PC1 and PC2 (bottom). (I) PCA of BPs from E12.5 to PN showing their transcriptional dynamics. (J) Heatmap of genes that have the highest contribution to the PC1 and PC2 for BPs, sorted by their weights (250 genes from each side). (K) PCA plots with projected genes shown as vectors illustrating their contribution. (L) Illustrating the trends (top), based on the position of samples along PC1 and PC2, and the gene expression profiles of top three genes from each side of PC1 and PC2 for BPs (bottom). (M) PCA of NBNs from E15.5 to PN showing their transcriptional dynamics. (N) Heatmap of genes that have the highest contribution to the PC1 and PC2 for NBNs, sorted by their weights (250 genes from each side). (O) PCA plots with projected genes shown as vectors illustrating their contribution. (P) Illustrating the trends (top), based on the position of samples along PC1 and PC2 and the gene expression profiles of top three genes from each side of PC1 and PC2 for NBNs (bottom). In D, H, L and P (bottom), the x-axis is embryonic days, and the y-axis is log2(TPM).

By contrast, *Neurod6, Cntn2*, *Slc17a7* and *Nfix* separated NSCs in the neurogenic phase along the PC2 negative axis from those in the expansion and gliogenic phases. Neurod6 is a Helix-Loop-Helix (HLH) transcription factors (TF) that plays a prominent role in neuronal differentiation (Sommer et al., 1996). *Neurod6* is transcriptionally activated by Neurog1 and Neurog2, two proneural HLH TFs that are downstream of Notch signaling (Ross et al., 2003). Neurod6 is associated with familial temporal lobe epilepsy and attention deficit-hyperactivity disorder in humans (Tutukova et al., 2021). Cntn2 is a member of the Contactin family of immunoglobulin cell adhesion molecules and functions in neuronal differentiation, determination, and migration as well as axon guidance (Mohebiany et al., 2014). *Cntn2* is located at 1q32.1, a region associated with microcephaly and mutations in *Cntn2* cause familial adult myoclonic epilepsy 5 (FAME5) (Mohebiany *et al*., 2014; Rickman et al., 2001; Stogmann et al., 2013). Slc17a7 is a transmembrane channel and urea transporter. It is selectively expressed in NSCs compared to BPs and NBNs and its expression increases with developmental stage. The function of Slc17a7 in NSCs remains to be shown. *Nfix* is a member of the nuclear I family of TFs. *Nfix* regulates NSC proliferation and differentiation both during embryonic development and in the adult and has been proposed to be a tumor suppressor in gliomas (Heng et al., 2015; Stringer et al., 2016). Loss of *Nfix* is associated with increased proliferation in the SVZ of the embryonic brain and delayed gliogenesis (Heng *et al*., 2015; Stringer *et al*., 2016). In summary, the unbiased analysis of gene expression revealed novel markers of stage specific NSCs and potential regulators of differentiation in the dramatic switch from the expansion to neurogenic phases of cortical development.

Conversely, *Pdgfra, Olig1, Gpr17, Tnc* and *Bcan* contributed prominently to the PC2 positive axis, segregating NSCs in the gliogenic phase of cortical development (Figure 2B- D). *Pdgfra, Olig1* and *Tnc* are known markers of glia cells and are upregulated in astrocytic and oligodendrocytic lineages (Di Bella *et al*., 2021). Tnc is a marker of outer radial glial cells during human brain development (Nowakowski *et al*., 2017). *Bcan* is expressed by immature oligodendrocytes and their precursors (Ogawa et al., 2001). *Bcan* and *Tnc* are implicated in the characteristic invasiveness of low-grade astrocytoma (Varga et al., 2012). Thus, the expression of these genes provides a distinctive signature for late phase NSCs during cortical development.

The unbiased computational approach identified genes important in NSC maintenance and differentiation as well as a plethora of novel and dynamically expressed NSC genes (Pollen *et al*., 2015; Telley *et al*., 2019a; Telley *et al*., 2016). We validated the expression of the novel signature genes predicted by the PC separations by RT-qPCR on independent biological replicates (Figures 2B and S2B, C; Table S2). We randomly selected genes differentially expressed by NSCs during the expansion, neurogenic and gliogenic phases. *Ccnd1, Crabp2, Hbb-bh1* are highly expressed during the expansion phase; *Bcl11b, Cntn2, Id2, Satb2* during neurogenic phase; and *ApoE, Aqp4, Sparcl1, Tril* during the gliogenic phase (Figure S2C and Table S2). By clustering the gene expression profiles we identified pools of genes that follow the same transcriptional trajectory in NSCs over time. This implied either co-regulation at the transcriptional level or gene expression associated with distinct cell states (Figure S2D-G). Some genes within these profiles are typical markers of the different phases of NSC development. For example, Neurog2 and Cspg4 mark the neurogenic and gliogenic phases of NSCs, respectively. Others, including Shh, mark specifically the early expansion phase and their expression is low upon the onset of fate determination (Figure S2D-G).

To address the transcriptional changes among the BPs and NBNs, we excluded the predominant variance resulting from the NSCs shifts in gene expression by computing PCs of all the samples orthogonal to the first two PCs of the NSCs. PCA of the remaining variables clustered all NSCs together indicating their underlying identity. BPs and NBNs separated in transcriptional space. Thus, the data orthogonal to the first PCs of NSCs enhanced the differences between NSCs, BPs and NBNs (Figures 2E) and increased the separation of BPs and NBNs, revealing a clear separation of BPs at early (E12.5-E14.5) and late (E15.5-PN) time points of cortical development (Figures 2E, F and S2H). In these analyses, NBNs showed less transcriptional dynamics over time. We performed pairwise comparisons to reveal DEGs genes contributing to PCs in the orthogonal analyses. Genes including *Dlx1, Dlx5* and *Dlx2* separated NSCs while *Tbr2*, *Nhlh1* represent the highest loading along the orthogonal PC1 negative axis. *Crym, Pf4* and *Crlf4* separated BPs and NBNs along the orthogonal PC1 positive axis (Figures 2F-H, Table S2).

To investigate the stage-correlated changes in gene expression by BPs and NBNs, we performed PCAs on BPs and NBNs separately. Despite being selected based on differences in level of *Tbr2::GFP* expression, PCA displayed continuous dynamics in these populations over time. However, the first PC was sufficient to separate BPs based on developmental stage (Figures 2I-L and S3A). From these analyses we identified *Fezf1, Samd3, Robo3* to be highest in early BPs while *Tac2, Dhrs3, Sh3rf3* were expressed higher by late BPs. Therefore, we could define distinct gene profiles for BPs that reflected their developmental stage implying that BPs are also a heterogeneous population of intermediate cells. Similarly, the first PC was also sufficient to separate NBNs over the course of development (Figures 2M- P, S3B). In order to validate the novel signature genes separating BPs and NBNs, we performed RT-qPCR on independent biological replicates (Figure S3C, D and Table S2) which confirmed the differential expression of the signature genes between early BPs, late BPs and their corresponding NBNs. *Cckar, Kif2c, Uncx, Robo3* were highly expressed by early BPs while *Loxl1, Unc5d, Ezr* were highly expressed by late BPs. On the contrary, NBNs displayed high expression of *Mef2c, Usp43, Lrfn5, Ntsr1* and *Gucy1a3* (Figure S2D). A more comprehensive list of these DEGs is available in Table 2. Thus, by our preliminary analyses of gene expression, we demonstrate dynamics in NSCs, BPs and NBNs and have identified novel signature genes which are binary and unique for these populations.

### Temporal dynamics in transcriptional landscapes of NSCs, BPs and NBNs is based on TF nodes and networks

To characterize the transcriptional states of NSCs, BPs and NBNs and map the activities of TFs throughout cortical development, we performed an Integrated System Motif Activity Response Analysis (ISMARA) (Artimo et al., 2016; Balwierz et al., 2014). ISMARA infers the regulatory state of samples as TF *‘motif activities’* (Figure 3A). PCA of the motif activities revealed maximum variance was dominated by NSCs, in agreement with the observations made based on mRNA expression (Figure 3B and 2A). Therefore, we split the data into two subsets, first to analyze the NSCs from all time points in isolation and second to analyze NSCs from the neurogenic phase compared to BPs and NBNs.

**Figure 3:**
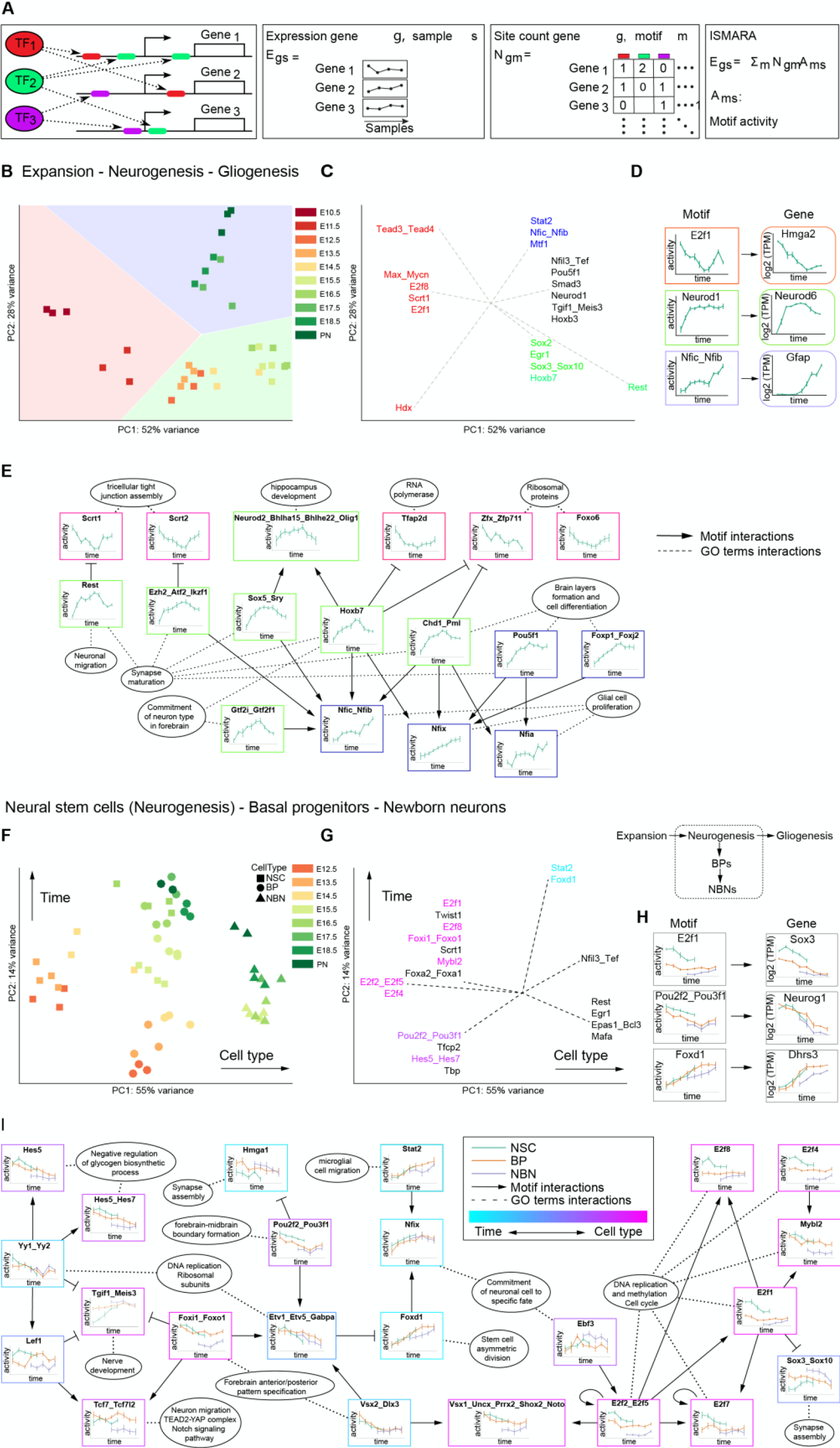
Dynamics of transcriptional network changes with ISMARA in different populations over time. (A) The ISMARA model’s promoter expression as a linear combination of the TF binding motifs activity that are present in the promoter region. (B) PCA on motif activity for NSCs for all time points, on the first two components, representing 80% of the total variance. The background color represents the three phases expansion (red, E10.5-E11.5), neurogenesis (green, E12.5-E15.5) and gliogenesis (purple, E16.5-PN). (C) Top 20 motifs contributing the most to the first two PCs, projected on the first two PCs. Each rectangular node of the network is a motif containing its activity plot in NSCs over time. The elliptical nodes represent the top GO categories. Each arrows represents gene activation while the stop lines represent gene repression. The dotted lines depict the main GO categories associated with the predicted targets of the connected motifs. The colors define the relative motif activity in the three phases, expansion (red), neurogenesis (green), gliogenesis (blue). (D) Examples of motifs (one per phase) regulating the genes identified from gene expression analyses, contributing highest to the PC1 and PC2 (figure 2 for NSCs). (E) Directed graphical representations of the main motif-motif interactions and the gene ontology and biological functions of the target genes. Each motif is shown with the color defining the zones of expansion, neurogenesis and gliogenesis along with plots of its activity. (F) PCA on motif activity for NSCs only from the neurogenesis phase, BPs and NBNs, representing 69% of the total variance. (G) Top motifs contributing the most to the first two principal components, projected on the first two principal components from (F). (H) Examples of motifs (one per cell type, NSCs- green, BPs- orange and NBNs- purple) regulating the genes identified from gene expression analyses, contributing highest to the PC1 and PC2. (I) Directed graphical representations of the main motif-motif interactions and the gene ontology and biological functions of the target genes. Each motif is shown with the color defining the cell types NSCs (green), BPs (orange) and NBNs (purple) along with plots of its activity.

The PCA for motif activity in NSCs showed that 80% of the variance was captured by the first two PCs, dividing the NSC samples into expansion, neurogenic and gliogenic phases (Figure 3B). We identified the top 20 TF binding motifs that contributed most to the variance in the PC1 and PC2 and displayed these as motif projections on the same subspace of gene expression (Figures 3C, S4A). Each motif is predicted to impinge on the expression of target genes although some TFs share highly related motifs (Artimo *et al*., 2016; Balwierz *et al*., 2014). For example, the TF E2f1 targets the promoter of *Hmga2*, Neurod1 targets *Neurod6* and Nfib (Nfic) targets *Gfap* (Figure 3D). Upon in-depth analyses of the TF motifs and their associated target genes, we observed a strong coherence with the transcripts identified in Figure 2B-D, thus validating the ISMARA approach.

ISMARA predicts the interactive regulatory networks of TFs in all cell types. Each edge of the network is characterized by the likelihood of an interaction. We selected the top motif-motif interactions to draw a simplified yet representative regulatory network in NSCs in the three phases of cortical development (Figure 3E). A large proportion of motifs cross-regulate each other and involve genes that are known to be involved in cortical development. Scrt1 and Scrt2 are active in NSC during the expansion, while Hoxb7 and Sox5 are active during the neurogenic phase, and Nfix, Nfia are involved in glial cell specification (Bel-Vialar et al., 2002; Lai et al., 2008; Paul et al., 2014; Zhou et al., 2015). Intriguingly, the active network motifs in the neurogenesis phase are predicted to repress the genes involved in expansion. Additionally, towards the end of neurogenesis, the activities of these motifs reduce and they in turn are predicted to activate genes to induce the transition to gliogenesis (Figure 3E). These analyses predict complex and dynamic, interconnected gene regulatory networks that can act independently but show a large degree of synergy.

We analyzed the TF motif dynamics of NSCs, BPs and NBNs together. Due to the dominant nature of the PC1 and PC2 of the NSCs motifs, we removed these for subsequent analyses and identified the top TF motifs determining the separation of NSCs, BPs and NBNs by PCA (Figure S4B, C). In accordance with our gene expression analyses, we performed parallel analyses for BPs and NBNs based on TF motif activities and found similarities in separation, identifying top selective nodes. These nodes are represented both as projections on the same subspace and as profiles of activities as determined from ISMARA (Figure S4D- I). To understand the relationship between the different cell types, we analyzed the dynamics in neurogenic NSC, BP, NBN TF activity over time. PC1 separated the three cell types into three clusters (Figure 3F). PC2 captured changes over time and surprisingly the dynamics in these motifs are shared by NSC, BP and NBNs (Figure 3F). We identified novel gene-sets defining the evolution of NSC, BP and NBNs over time (Figure 3G, H).

We identified the main interactions within the NSC, BP and NBNs and represent these as a predicted interconnected network (Figure 3I). A subnetwork of E2f family motifs (E2f1, E2f2, E2f4, E2f5, and E2f8) which are involved in DNA replication, methylation, and cell cycle, and positively impinge on Mybl2 (a cell cycle regulator) and paired-like homeodomain Vsx1-like TF activity and counteract Sox3/Sox10 activity. Although, these motifs characterize differences between cell types, they remain relatively constant over time. We observe TF motifs that show similar activities across cell-types but increase in their activity over time. Foxd1 and Stat2 are examples of motifs regulating neural and glial development through Nfix (Figure 3I). Conversely, some TF motifs show cell-type specificity (Hes1, Hes5, Meis3, Tcf7 and Foxo1). Interestingly, Hes1, Hes5, and Tcf7 are all primary regulators of Notch signaling in NSCs (Figure 3I). From the ISMARA predictions, interactions of motifs highly active in NSCs during neurogenesis project towards motifs that are highly active in other cell types and phases. This directionality indicates strong intrinsic properties of neurogenic NSCs.

### Single-cell RNA sequencing reveals underlying heterogeneity in NSCs, BPs and NBNs

PCA at the population level revealed extensive changes in transcriptome in NSCs, BPs and NBNs over time. We addressed heterogeneity within each cell population by analyzing the transcriptional landscapes at the single-cell level by single cell RNA-Seq (scRNA-Seq) (Figure S5A). The single-cell transcriptomes of highly variable genes (HVGs) revealed a low heterogeneity within the NSCs during expansion and gliogenesis (Figure 4A). By contrast NSCs during the neurogenesis phase (E13.5 and E15.5) were heterogeneous (Figure 4A). To validate that the scRNA-Seq data were representative of the population data, we averaged the single-cell transcriptomes of a specific time point and projected them on the PC matrices of the population samples. The averaged single-cell data superimposed on the population samples and followed the same transcriptional trajectory over time and confirmed that the single cell transcriptomes reflected the heterogeneity of the population at the respective time point (Figure 4B). Therefore, the single cell heterogeneity in NSCs during cortical development is representative of the biological changes in single NSC gene expression over time.

**Figure 4:**
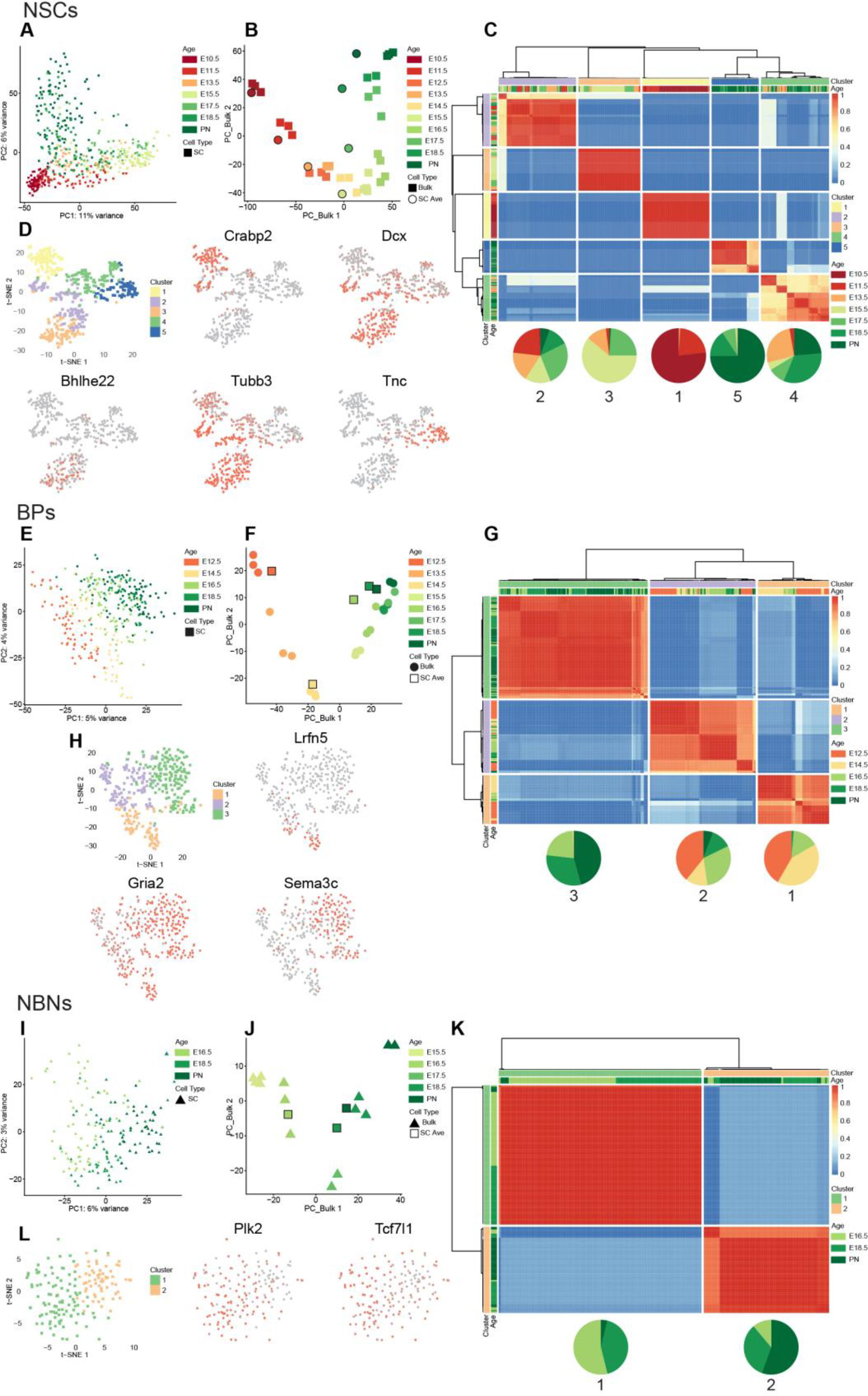
Heterogeneity of NSCs, BPs and NBNs at single cell level. (A) PCA of NSC single cells, using the top 2000 highly variable genes obtained from bulk NSCs. (B) Projection of average single cells of NSCs at each time point on the first two PCs of bulk NSCs using the top 2000 highly variable genes obtained from bulk NSCs. (C) Clustering of assignment matrix of NSC single cells using k-means and hierarchical clustering. (D) Marker genes that are up/down regulated in each cluster of NSCs. (E) PCA of BP single cells, using the top 2000 highly variable genes obtained from bulk BPs. (F) Projection of average BP single cells on the first two PCs of bulk BPs using the top 2000 highly variable genes obtained from bulk BPs. (G) Clustering of assignment matrix of NBN single cells using k-means and hierarchical clustering. (H) Marker genes that are up/down regulated in each cluster of NBNs. (I) PCA of NBN single cells, using the top 2000 highly variable genes obtained from bulk NBNs. (J) Projection of average single cells of NBNs on the first two PCs of bulk NBNs using the top 2000 highly variable genes obtained from bulk NBNs. (K) Clustering of assignment matrix of NBN single cells using k-means and hierarchical clustering. In C, G and K, heatmaps represent the hierarchal clustering of assignment matrix of single cells after 500 times applying k-means clustering. The optimal number of clusters is selected based on the Silhouette coefficient. It is “1” (red) when two cells are always clustered together, “0” (blue) when two cells never fall in the same cluster. Pie charts represent the percentage of single cells at each time point in each cluster.

k-means clustering divided the NSCs into five cell clusters and revealed DEGs across these clusters (Figures 4C, S5C-F and Table 4). The five NSC types were unequally represented over time. NSC type 1 (cluster 1) were present almost exclusively at E10.5 and E11.5 and represent the major NSC transcriptional status in the expansion phase. NSC cluster 5 was the predominant NSC type during the later, gliogenic phases of corticogenesis. NSC clusters 2-4 were found during multiple phases of development from E11.5 and expansion through neurogenesis to gliogenesis (Figures 4C).

Visualization of the single cell data by t-SNE also showed separation of the five NSC cell-types (clusters 1-5: Figure 4D). Projection of gene expression onto the t-SNE identified cluster specific expression. *Crabp2* and *Tnc* marked clusters 1 and 5, respectively while *Tubb3* and *Dcx* were expressed in a more expanded domain across multiple NSC clusters (Figure 4D). The heatmap shows a more comprehensive list of distinct signature genes for the five clusters (Figure S5F and Table 4). The GO analyses and process networks for gene expression by the NSC clusters is shown in Table 5.

We analyzed heterogeneity within the BPs from E12.5-PN1 by scRNA-Seq. BPs showed an age-related difference is gene expression with heterogeneity distributed along the PC2 (Figure 4E). We pooled the single cell sequences at each time point and plotted these averaged values on a PCA defined by the HVGs identified from the BP analysis at the population level (Figure 4F). Strikingly, the averaged single-cell data superimposed on the biological replicates at population level supporting that the individual BP scRNA-Seq data are representative of the populations (Figure 4F). The analysis indicated a distinct, time-dependent dynamic in gene expression from E12.5-PN1. Clustering the single BPs based on k-means revealed three distinct clusters (Figures 4G, S5G-I and Table 4). Cluster cell-type 3 cells are present mostly at later developmental stages (E16.5-PN1). Conversely, cluster 1 and 2 cells were more prominent at earlier times (Figures 4G). We visualized the BP clusters by t-SNE and plotted some signature genes for each cluster (Figures 4H, S5J and Table 4). We performed similar analyses for NBNs expecting a large heterogeneity over time due to the plethora of neuron types generated in the dorsal cortex. scRNA-Seq confirmed a broad spread in the NBNs with a component of time over the PC1 (Figure 4I).

We pooled the single cell NBN data at each time point and plotted these averaged values on a PCA defined by the HVGs from the NBN analysis at the population level (Figure 4J). Both the population and averaged single cell samples followed a time-dependent trajectory in gene expression consistent with the sequential generation of neurons forming the deep and superficial layers of the isocortex (Figure 4J). k-means clustering revealed the reduced heterogeneity in the NBNs compared to NSCs and BPs and identified 2 major clusters (Figure 4K). Cell belonging to cluster type 1 NBNs were present at earlier time points (E15.5 and E18.5), while cells belonging to cluster type 2 were underrepresented at E15.5 and were mostly present at PN1 (Figure 4K and Table 4).

t-SNE representation of the NBNs belonging to the two cell clusters showed the sparse distribution of type 1 cells reflecting the single cell heterogeneity within this cluster. Type 2 NBNs clustered more tightly than cluster 1 cells. Due to this heterogeneity, it was difficult to pinpoint single gene signatures for the two NBN clusters (Figure 4L, S5N and Table 4). These findings demonstrate an unprecedented heterogeneity in NSCs, BPs and NBNs over time and a dynamic shift in gene expression of these cells at the population and single-cell levels. The scRNA-Seq data enabled a high-resolution definition of gene signatures for each cluster (cell type) of NSCs, BPs and NBNs.

### Neuronal specification markers show sequential waves in gene expression and massive heterogeneity at the single cell level

During cortical development, morphologically and physiologically unique classes of neurons are formed sequentially in waves throughout neurogenesis (Greig *et al*., 2013; Molyneaux *et al*., 2007; Telley *et al*., 2016) (Figure 5A). Several gene combinations have been identified that classify the distinct subtypes of projection neurons in the cortex (Greig *et al*., 2013; Molyneaux *et al*., 2007). We selected and curated an extensive list of known patterning and neuronal subtype marker genes from the literature and analyzed their expression dynamics in the NSC, BP and NBNs at the population and single cell levels.

**Figure 5:**
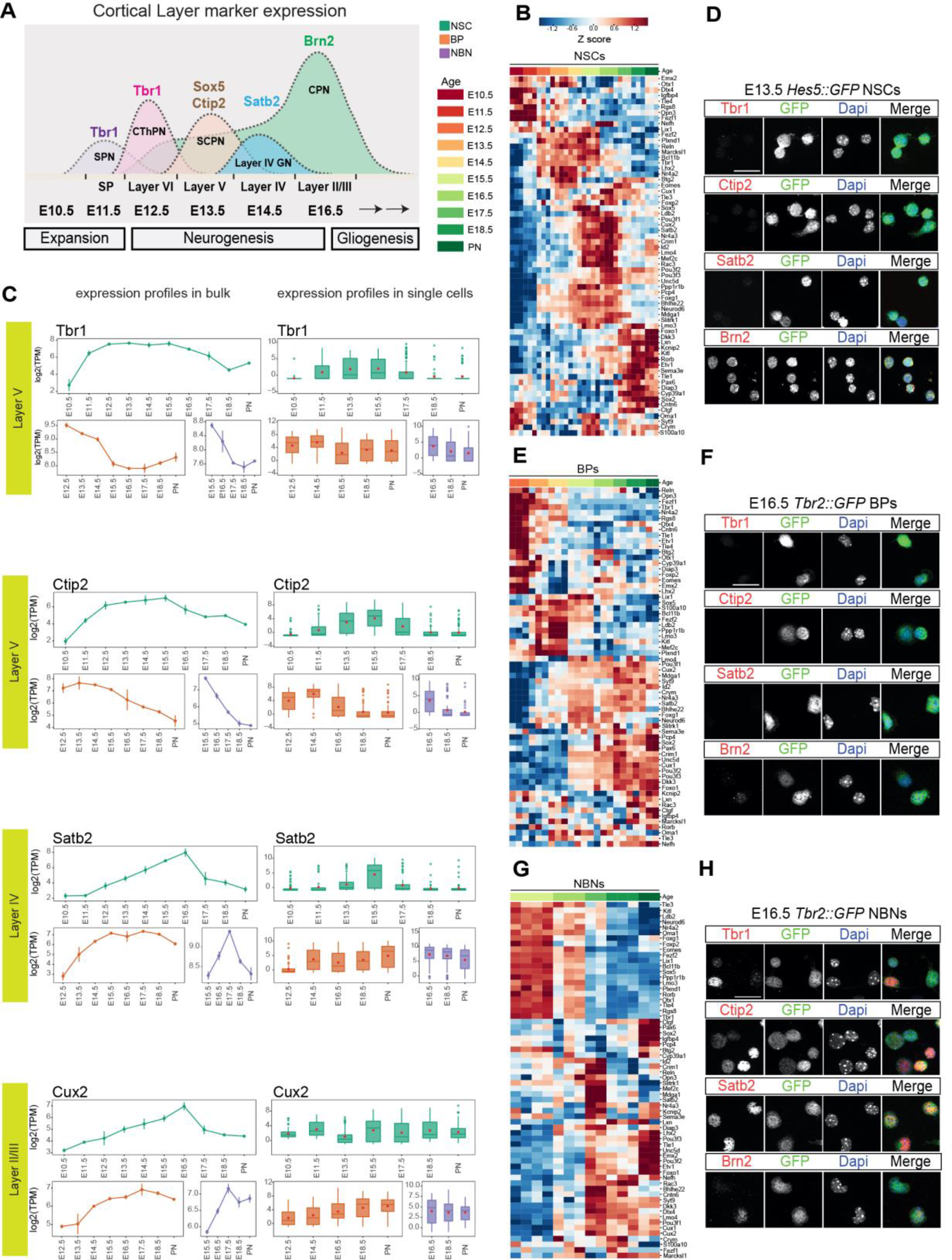
Dynamic expression of neuronal specification factors in NSCs, BPs and NBNs. (A) Illustration of distinct projection neurons born sequentially during the course of neurogenesis. (B) Heatmap illustrating the dynamics of expression of cortical layering markers in NSCs at population level. (C) Examples of expression dynamics of deep layer markers Tbr1, Ctip2 and upper layer markers Satb2, Cux2 in NSCs, BPs and NBNs, profiles at population level (left) and single cell level (right). (D) Experimental validation of NSCs isolated at E13.5 using *Hes5::GFP* transgenic embryos, showing no detectable protein for Tbr1, Ctip2 and Satb2. NSCs do express Brn2(Pou3f2) *in vitro* and *in vivo* at protein level. (E) Heatmap illustrating the dynamics of expression of cortical layering markers in BPs at population level. (F) Experimental validation of BPs isolated at E16.5 using *Tbr2::GFP* transgenic embryos, showing no detectable protein for Tbr1, Ctip2 and Satb2. (G) Heatmap illustrating the dynamics of expression of cortical layering markers in NBNs at population level. (H) Experimental validation of NBNs isolated at E16.5 using *Tbr2::GFP* transgenic embryos, showing protein expression for Tbr1, Ctip2, Satb2 and Brn2(Pou3f2). In B, E and F, heatmaps are based on z-score of log2(TPM) expression values.

Surprisingly, transcription of the neuronal subtype genes showed sequential and developmental wave-like patterns of expression even in NSCs, at the population level (Figure 5B, S6A). At the single cell level, these developmental waves were recapitulated in NSCs, albeit with a pronounced heterogeneity at each time point (Figures 5C). Particularly those genes commonly used to define neuronal subtypes and cortical layers later in development (*Tbr1* and *Ctip2* - Layers V and VI; *Satb2* and *Cux2* - Layers IV and II/III) showed characteristic and transient dynamics in expression one to two days prior to the established birth-date of the neurons (Molyneaux *et al*., 2007). We plated the freshly FACsorted *Hes5::GFP* positive NSCs but could not detect expression of these neuronal specification factors at the protein level by immunocytochemistry (Figure 5D). This suggested that the transcriptional program that defines cortical neuron subtypes is initiated in NSCs long before their exit from cell cycle.

We performed similar computational analyses on the BPs and NBNs and identified similar sequential waves of cortical neuron gene expression correlating with the birth date of the respective neuron subtype (Figures 5C, E, F and S6B). Similarly, we could not detect protein expression in the BPs acutely isolated form the developing cortex (Figures 5C, 5E, 5F and S6B). However, and as expected, NBNs expressed proteins associated with neuron subtypes of definitive cortical layers (Figures 5C, 5G, 5H and S6C).

These striking findings indicates that neuronal specification programs start early in the lineage, in NSCs and BPs, and continue into the NBNs. At the single-cell level, some E10.5 NSCs expressed high levels of deep layer neuronal markers including Cux2 while later NSCs expressed both deep and upper-layer neuronal markers (Figure 5C, S6D). This explains the seemingly controversial Cux2 lineage tracing experiments described previously (Franco *et al*., 2012).

### Signaling pathway effectors show dynamic expression in the neurogenic lineage

Signaling pathways impinge on downstream effectors to regulate NSC fate decisions during corticogenesis. The crosstalk between signaling pathways and the integration of their target effectors governs stem cell maintenance and fate. However, it remains unclear to which signals NSC, BPs and NBNs are competent to respond *in vivo*. In order to evaluate susceptibility to paracrine signaling molecules and the dynamics in this responsiveness, we selected those genes designated to be receptors in the databases and analyzed the expression of the 440 receptors that showed variable gene expression throughout the neurogenic lineage. The resulting extensive gene expression profiles could be divided into three groups (Figure 6A, B):

1. Receptors that are highly expressed by NSCs through the most of cortical developmental (Figure 6A, B). These receptors, including those for Wnt (*Fzd5, 7, 9*), Notch (*Notch1, 2, 3*), Fgf (*Fgfr2, Fgfr3*) and Shh signaling (*Smo, Ptch1*), are part of pathways involved in stem cell maintenance (Blaschuk and ffrench-Constant, 1998; Bray, 2006; Fukuchi-Shimogori and Grove, 2001; Gaiano and Fishell, 2002; Imayoshi *et al*., 2010; Itoh and Ornitz, 2004; Iwata and Hevner, 2009; Louvi and Artavanis-Tsakonas, 2006; Rash et al., 2011; Sahara and O’Leary, 2009; Shimojo et al., 2008; 2011; Wang et al., 2011).
2. Receptors that are highly expressed by NSCs during neurogenesis and later stages and by BPs and NBNs which we refer to as neurogenic (Figure 6A).
3. Receptors that are highly expressed predominantly at later stages of development in the NSCs during the gliogenic phase. These we refer to as gliogenic pathways and include the receptors for known ligands involved in gliogenesis, including Tgf-beta/BMP signaling (*Tgfbr2, Bmpr1a, Bmpr1b*) and Il6/Lif signaling (*Lifr, Il6st*) (Ebendal et al., 1998; Gomes et al., 2005; Pollen *et al*., 2015; Rodriguez-Martinez and Velasco, 2012).

**Figure 6:**
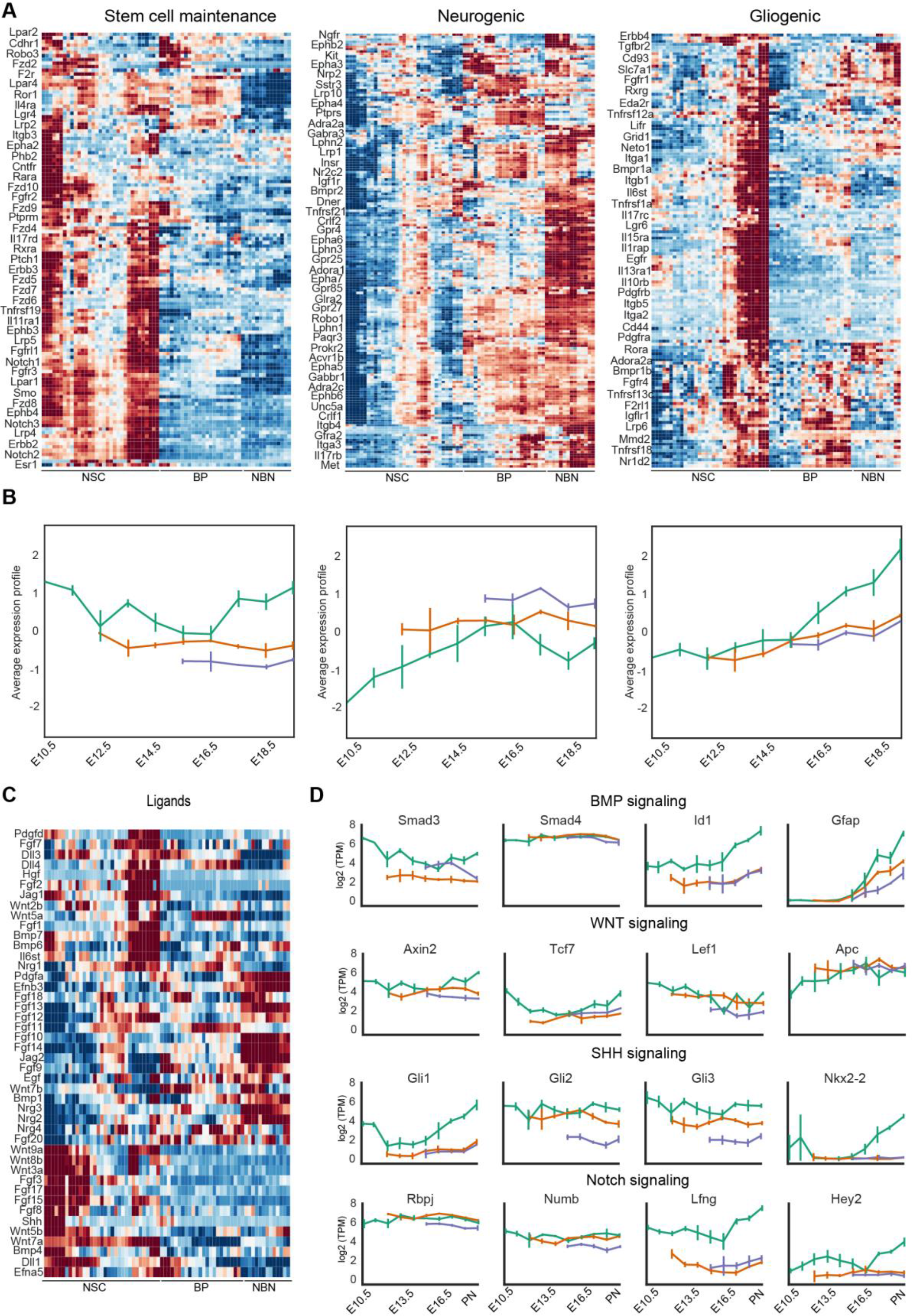
Dynamic expression profile of signaling receptors during corticogenesis. (A) Heatmaps representing the expression profile of signaling receptors that can be divided into three main groups based on k-means clustering of z-scored log2(TPM) expression values: stem cell maintenance (121 receptors), neurogenic (180 receptors) and gliogenic (139 receptors). Names of selected receptors are displayed. For the complete list please see Supplementary Information. Expression profiles are represented by their z-score. (B) Average expression profile of each cluster for NSCs (green), BP (orange) and NBN (purple). Solid line represents the average z-score, while the area represents the standard deviation estimated from different biological samples. (C) Heatmap representing the expression profile of ligands from selected signaling pathways, based on the z-scored log2(TPM) expression values. (D) Expression profile of selected target or modulator of key signaling pathways: BMP, Wnt, Shh and Notch signaling.

The neurogenic niche during corticogenesis provides local autocrine and paracrine signals but also responds to blood-born ligands and factors in the fluid of the telencephalic vesicles. We assessed the potential local signals in the NSCs, BPs and NBNs by examining the expression of ligands for the top, regulated receptors (Figure 5C). Similar to the expression profile of their cognate receptors, the expression of some ligands could be divided into three clusters. Ligands expressed predominantly by NSCs during the expansion phase of corticogenesis, ligands expressed predominantly by NSCs in the gliogenic phase, and ligands expressed mostly by neurons that act as paracrine signals back to the progenitors. Many Wnt ligands and their receptors are expressed by NSCs suggesting autocrine signaling. One notable exception being *Wnt7b* which is prominently expressed by NBNs and its canonical receptor Fzd7 also by NSCs at the expansion and gliogenesis phases. By contrast, although their cognate receptors were mostly expressed by NSCs, the Fgf ligands were divided into two major groups, those expressed mainly by NBNs, and those expressed mainly by NSCs (Figure 6A, C).

As a proof of concept, we also evaluated selected modulators and effector targets of some of the key signaling pathways including Bmp, Wnt, Notch and Shh signaling (Figure 6D). The expression of many target genes of these pathways reflected the expression of their respective receptors suggesting that not only are the ligands available and the receptors expressed but the pathways may be active throughout the neurogenic lineage (Figure 6D).

### bHLH TFs are dynamically and heterogeneously expressed by NSCs

bHLH TFs are notably involved in the control of neurogenesis and brain development downstream of many pathways including Notch, BMP, TGFβ and Wnts. We analyzed the expression profile of the bHLH family genes. The bHLH TFs could be grouped into three classes based on their expression profiles:

1. bHLH factors related to NSCs maintenance, including *Hes1, Hes5, Hey1* and *Id4*, are highly expressed by NSCs (Figure 7A, B).
2. bHLH factors related to neuronal commitment and differentiation. For example, the proneural differentiation bHLH genes including *Neurog2, Neurod2* and *Neurod6*, which are expressed by NSCs during the neurogenic phase and by BPs, but their expression is lower in NBNs (Figure 7A, C).
3. bHLH genes with expression associated prominently with gliogenesis and which are expressed at low levels by BPs and NBNs including *Olig1, Olig2* and *Id1* (Figure 7A, D).

**Figure 7:**
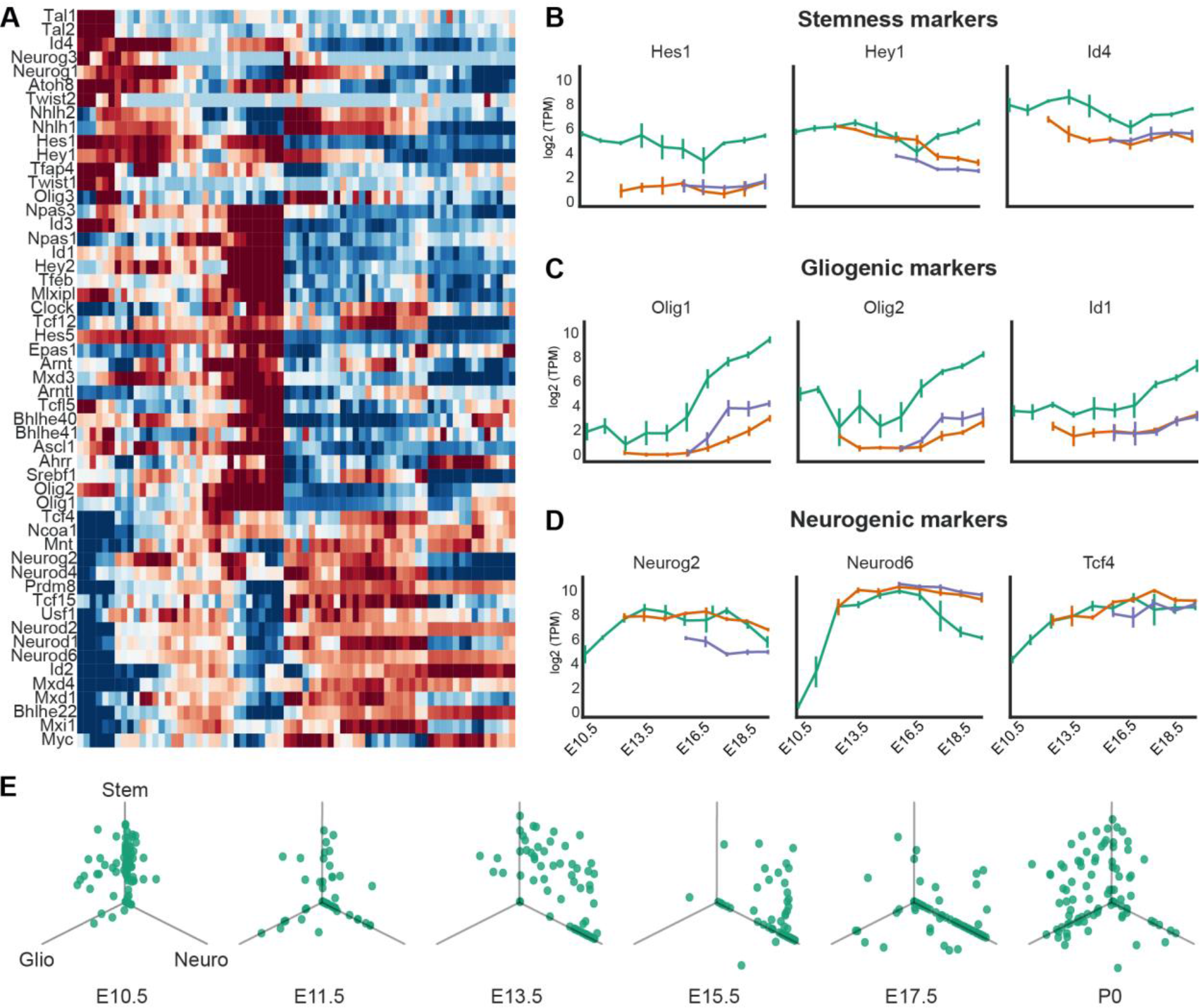
Dynamic and heterogenic expression profile of bHLH factors during forebrain development. (A) Heatmaps representing the expression profile of bHLH factors. Three main groups are observed based on k-means clustering of z-scored log2(TPM) of expression value: stem cell maintenance (high expression in the NSCs at early embryonic times and low in BPs and NBNs), neurogenic (high expression in the NSCs during neurogenesis and high expression in BPs and NBNs) and gliogenic (high expression in the NSCs at late embryonic times and low in BPs and NBNs). Expression profiles are represented by their z-score. (B) Expression profile of selected stem cell maintenance markers Hes1, Hey1 and Id4. (C) Expression profile of selected neurogenic markers Neurog2, Neurod2, and Neurod6. (D) Expression profile of selected gliogenic markers Olig1, Olig2 and Id1. (E) Expression of stem cell markers (Hes1, Hey1 and Id4), neurogenic markers (Neurog2, Neurod2, and Neurod6) and gliogenic markers (Olig1,2 and Id1) in NSCs during different embryonic time points in the single-cell levels. Each point represents the expression value of one single cell in log2(TPM).

We also identified a group of bHLH TFs expressed moderately by NSCs during the neurogenic phase of corticogenesis, but which are expressed by BPs and NBNs suggesting a role in neural commitment and differentiation (Figure 7C, 7D). At the single cell level, expression of the bHLH factors by the NSCs was highly heterogeneous, even at the same embryonic time point (Figure 7E). As expected at E10.5 most NSCs expressed high levels stemness markers (*Hes1, Hey1,* and *Id4)*, and low or no neurogenic associated (*Neurog2, Neurod2,* and *Neurod6*) and gliogenic associated bHLH TFs (*Olig1, Olig2,* and *Id1*). As neurogenesis initiated at E13.5, more cells started to express neurogenic markers and the number of cells expressing *Hes1, Hey1,* and *Id4* reduced but very few cells were expressing the gliogenic bHLHs. At E13.5 two major NSC populations were evident on the basis of bHLH expression, one expressing high stemness markers and low neurogenic markers, and another expressing low stemness markers and high neurogenic markers. However, there was also a subpopulation of cells that expressed both maintenance and neurogenesis associated bHLH TFs. One explanation for this populations is the oscillatory expression of stemness factors *Hes1* and *Hes5*, and their repression of the neurogenic targets including *Neurog2*. At later stages, when NSCs exit the neurogenic phase and enter gliogenesis (E15.5.-E17.5), the proportion of NSCs expressing the neurogenic bHLHs rather than the stem cell maintenance associated TFs increased. As the NSCs transitioned into gliogenesis and towards PN1, the proportion of NSCs expressing *Neurog2, Neurod2,* and *Neurod6* diminished with a concomitant increase in gliogenic factor (*Olig1, Olig2,* and *Id1*) expressing cells (Figure 7E). Strikingly, some NSCs in the gliogenic state coexpressed the maintenance bHLH TFs. This confirms previous observations that Notch signaling, and the expression of Hes-related TFs is linked to glial commitment of NSCs.

## Discussion

The temporal dynamics in gene expression during lineage commitment throughout ontogeny of the cerebral cortex remains unclear. Advances in scRNA-Seq and gene cluster analysis have given unprecedented insights into cellular heterogeneity in the mammalian cortex, also in fetal humans. Particularly when trying to understand the genetic regulation of cell diversification from stem and progenitors, these snapshots of cellular transcriptomics are used to define cellular state and therefore predict potential. One challenge for transcriptome analysis is to predict not only the future of a particular cell and its offspring but also its history. This is particularly confounded by highly dynamic gene expression over time-windows ranging from days to minutes.

Here we posed the questions of how gene expression changes in stem cells, progenitors, and newly formed neurons over time and whether we can uncover distinct traits and patterns within specific cell-types defined based not on an *ad hoc* identification using a selection of RNA transcripts but on functional properties. We characterized gene expression of the dorsal cortical neural lineages over time, focusing on the phases of expansion, neurogenesis and gliogenesis and on NSC, BP and NBN populations with definitive characteristics. We have created an extensive resource of dorsal cortical ontogeny which can be mined through an interactive web-based browser (Figure S7; http://neurostemx.ethz.ch/).

Development of the cerebral cortex is a dynamic process, however, and remarkably, our understanding of the lineage heterogeneity and changes in gene expression that accompany the formation of the different neuron subtypes and subsequent cortical layers is limited. The most widely accepted model of cortical development utilizes a common multipotent progenitor, which becomes progressively restricted in its fate over time. Unbiased computational analysis of our data revealed distinctive, stage-specific changes in gene expression not only in NSC, but also BPs and NBNs. These shifts in transcriptional space at the population and single cell levels reveal a heterogeneity in each of these cell populations and establish novel gene signatures defining five NSC, three BP and two NBN types. Remarkably, we show the presence of different NSC, BP and NBN types at the same developmental stage, and that these constitute different proportions of the particular populations at each stage. Although these findings do not disprove a common progenitor mechanism, they imply that the populations of NSC, BPs, and NBNs at any point in developmental time are composed of different proportions of cells with distinct transcriptomes which can be predicted by a panel of signature genes.

Our analyses of cells-types with definitive characteristics exemplifies potential dangers in a priori allocation of cell-type based on comparative gene expression and limited gene sets. For example, the transcriptomes of NSC in the neurogenic phase of cortical development are closer to BPs at this stage than they are to NSCs at E10.5 (expansion phase) or PN (gliogenic phase). Only by separating the NSCs, BPs and NBNs and analyzing their transcriptomes in isolation, was it possible to uncover specific signatures and determine cellular heterogeneity. The conjunction of gene expression dynamics and predicted transcriptional networks by ISMARA, identified active TFs and nodes in NSCs, BPs and NBNs over time. These TF motifs and activities also revealed the same directional trajectory and pathway of each cell type through transcriptional space as predicted by the mRNA expression. As ISMARA predicts the activity of more than 800 TFs and their targets in all cell types this data-set and resource will be valuable to explore and extrapolate the known regulatory networks to the missing novel nodes. Recently, we validated the Tead TFs in cortical development and elucidated different functions of Tead factors in NSCs (Mukhtar *et al*., 2020). The dynamic changes in the TFs in NSCs for example, reflects the sequential changes these cells undergo during corticogenesis. Our analyses determining the relationship among the neuronal lineage demonstrate a naturally occurring directionality, indicating strong intrinsic control. Moreover, among all the cell types NSCs seem to be most dynamic, be it at gene expression level, or the level of transcriptional networks. The NSCs follow a continuous path through the three phases of corticogenesis, supporting the neuronal origin from ‘common progenitors’, sequentially changing in transcriptional space. To this end, we identified signature genes of NSCs in expansion, neurogenic and gliogenic phases. As all the known genes depicted the dynamics of expression as expected, we believe that we provide more extensive lists of novel signature genes which could be used to identify NSCs in different phases. These signatures are like a ‘scorecard’ for the NSC population undergoing corticogenesis, some of which we have also validated experimentally (Figure S2B, C and S3C, D). Similar analyses for BPs and NBNs have yielded key signature markers which hold promise for further biological exploration. The up and down regulations of genes could be presumed to be the result of active or inactive downstream programs in these cells and their progeny. It is crucial to differentiate between early and late BPs, or different NBN populations across time in order to consolidate our knowledge about their downstream fate and function.

The microenvironment of the cells plays critical roles in regulating cell fate choices. We used these resources and explored some of the signaling pathways defining logic in NSC differentiation using a high-throughput microfluidic approach (Zhang et al., 2019). This validation was the tip of the iceberg and together with the recent developments in the field, we provide a consolidated resource, a comprehensive and systematic characterization of major progenitor pools in cortical development. Further biological validations of our predicted signaling and transcriptional nodes will provide more promise towards the deeper exploration of mechanisms controlling corticogenesis.

## AUTHOR CONTRIBUTIONS

V.T ., E.v.N, D.I., C.B., S.A., conceived and designed the project. T.M., K.E., C.B. and V.T. designed and performed experiments, evaluated, and interpreted the data. J.B., M.B., Z.K., P.G., A.G., analyzed the data. Z.K., S.V., M.O., R.S. and M.P. designed the website. T.M., J.B., M.B., Z.K., P.G, A.G., and V.T. wrote the paper and prepared the figures. All authors edited and proofread the manuscript.

## ACKNOWLEDGEMENTS

We thank the members of the V.T. laboratory for helpful discussions and Frank Sager for excellent technical assistance. We thank the Animal Core Facility of the University of Basel. This work was a part of a SystemX consortium- NeuroStemX.

## Supplementary Figures

**Supplementary figure 1:**
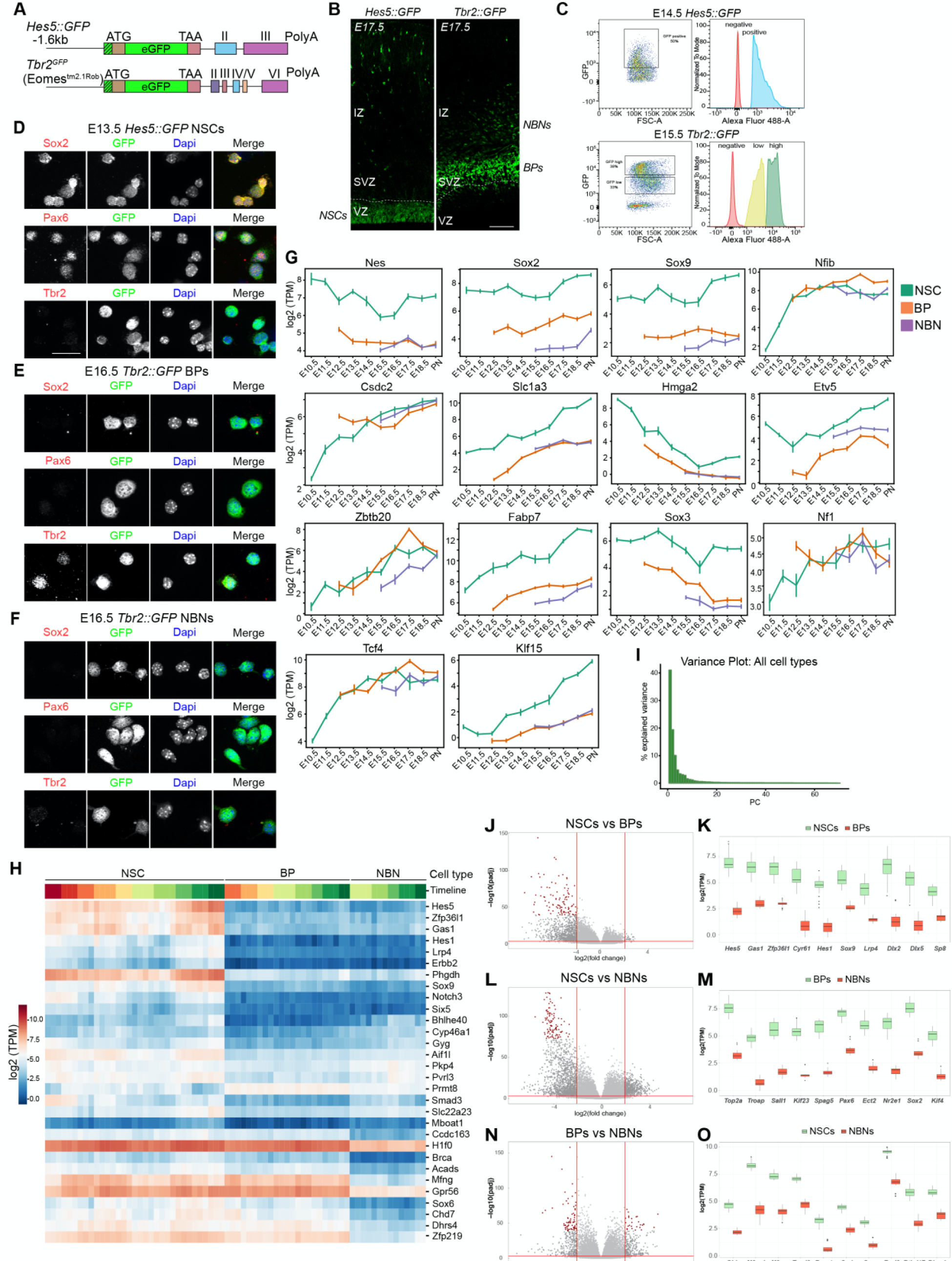
(A) Hes5::GFP and Tbr2::GFP transgenic mice used for cell isolation. (B) Expression of *Hes5::GFP* and *Tbr2::GFP* embryonic cortices at E17.5. Scale bar = 100μm. (C) Examples of FACS plots for GFP positive cell sorting at E14.5 *Hes5::GFP* and E15.5 *Tbr2::GFP*. (D-F) Expression validation of *Hes5::GFP* and *Tbr2::GFP* positive cells after FAC sorting *in vitro*. Scale bar = 20μm (G) Expression plots of some known markers of NSCs. (H) Heatmap showing differentially expressed genes in three cell populations illustrating NSCs, BPs and NBNs vary in expression, based on z-scored log2(TPM) expression values. (I) Bar plot representing the proportion of variance covered by each PC in PCA of all cell types. (J, L, N) Volcano plots for DEG analysis for NSCs versus BPs, NSCs versus NBNs and BPs versus NBNs, respectively. Significantly DEGs are colored as grey and top 100 DEGs are colored by red. (K, M, O) Top ten DEGs for NSCs versus BPs, NSCs versus NBNs and BPs versus NBNs, respectively. (J-O) are related to analysis of Figure 1E. The range of p-values is very different: NSC (0.01%-0.4%), BP (1.6% – 4.9%), NBN (0.06%-0.2%). There are no good marker genes for BPs as their gene expression tends to be similar to either NSC or NBN.

**Supplementary figure 2:**
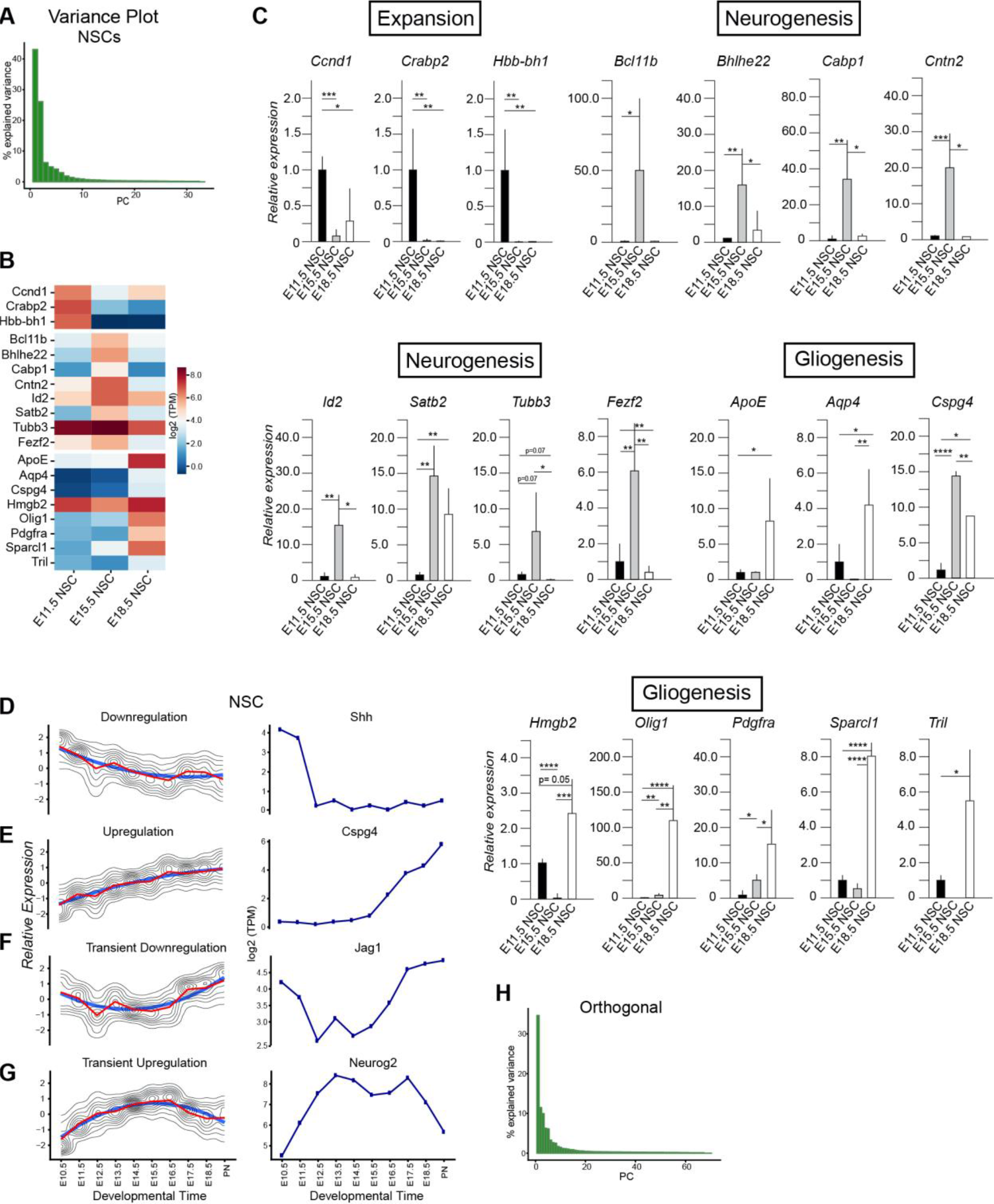
(A) Bar plot representing the variance coverage by PC corresponding to PCA plot in Figure 2(A). (B) Heatmap illustrating the expression changes in signature genes in time points corresponding to expansion, neurogenesis and gliogenesis. (C) qPCR validation of signature genes in three zones. Each time point has samples varying from N =3 to N= 7. (D-G) k-means clustering of z-scored log2 (TPM) gene expression profiles over developmental time course in NSCs with genes showing upregulation, e.g., Cspg4, downregulation, e.g., Shh, transient downregulation, e.g., Jag1, transient upregulation, e.g., Neurog2. (H) Bar plot representing the variance coverage by PC corresponding to PCA plot in Figure 2 (E).

**Supplementary figure 3:**
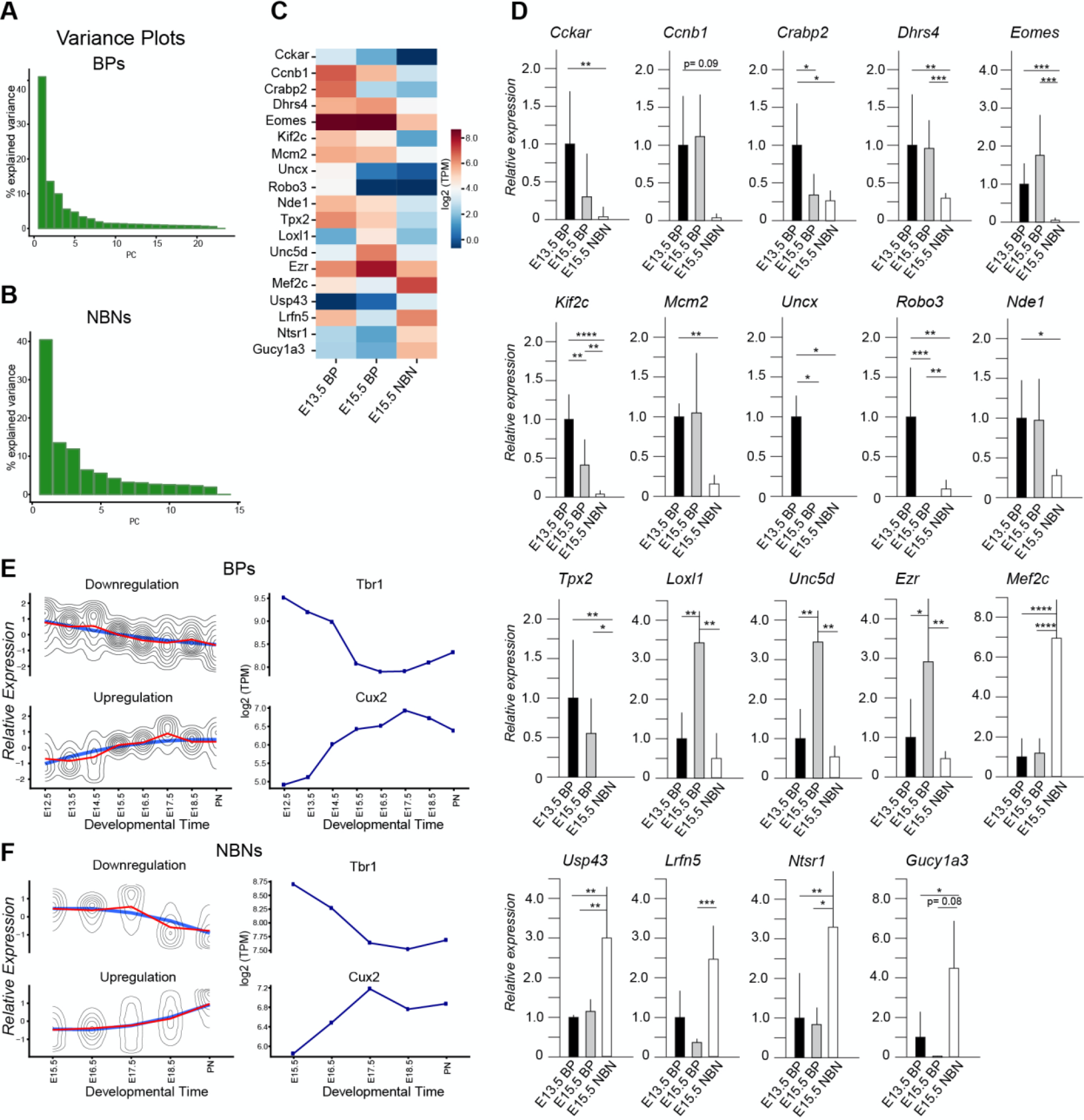
(A, B) Bar plots representing the variance coverage by PCs corresponding to PCA plot in Figure 2 (I, M). (C) Heatmap illustrating the expression changes in signature genes in time points corresponding to early BPs, mid-BPs and NBNs. (D) qPCR validation of signature genes for three sample types. Each time point has samples varying from N =3 to N= 7. (E) k-means clustering of z-scored log2(TPM) gene expression profiles over developmental time course in BPs with genes showing downregulation, e.g., Tbr1 and upregulation, e.g., Cux2. (F) k-means clustering of z-scored log2(TPM) gene expression profiles over developmental time course in NBNs with genes showing downregulation, e.g., Tbr1 and upregulation e.g., Cux2.

**Supplementary figure 4:**
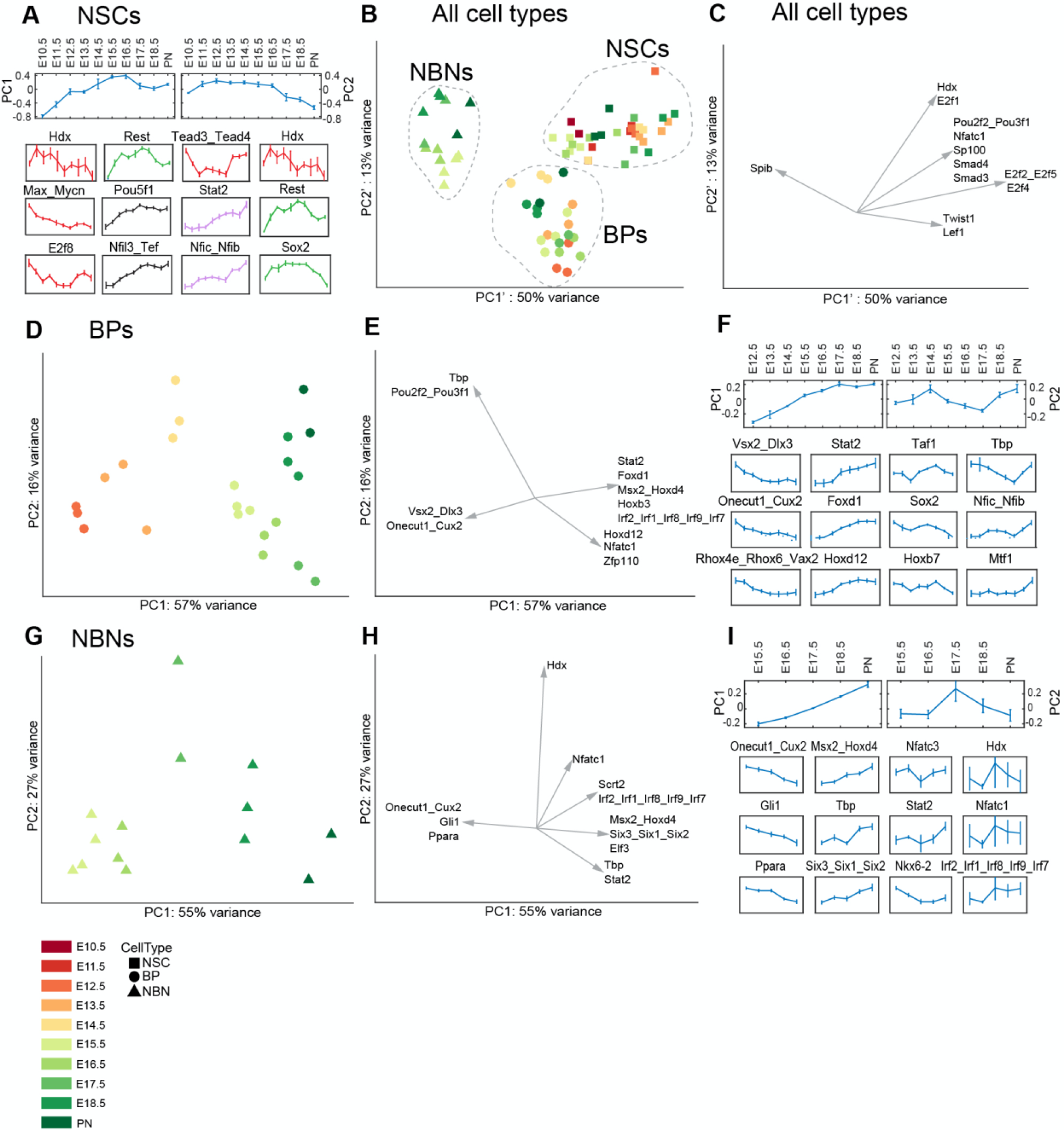
(A) Examples of dynamic motifs based on the PCA of NSCs (Figure 3B). Plots show the replicate average of samples across the sampling time for the first two principal components separately, and for the three motifs contributing the most to the first and second principal component, positively and negatively, separately. (B) PCA plot for all cell types (NSCs, BPs and NBNs) after removing the first two components of NSC variance (from Figure 3B). (C) Plot showing the projections of each cell type sample on the replicate average of motif activity, representing 63% of the total variance. (D) PCA analysis on motif activity for BPs for all time points, on the first two components, representing 73% of the total variance. (E) Top 12 motifs contributing the most to the first two principal components, projected on the first two principal components. (F) Examples of dynamic motifs based on the PCA of BPs. Plots show the replicate average of samples across the sampling time for the first two principal components separately, and for the three motifs contributing the most to the first and second principal component, positively and negatively, separately. (G) PCA on motif activity for NBNs for all time points, on the first two components, representing 82% of the total variance. (H) Plot showing the projections of each cell type sample on the replicate average of motif activity, representing 82% of the total variance. (I) Examples of dynamic motifs based on the PCA of NBNs. Plots show the replicate average of samples across the sampling time for the first two principal components separately, and for the three motifs contributing the most to the first and second principal component, positively and negatively, separately. In A, F and I bottom, the y-axis is the embryonic day and x-axis is log2(TPM) expression values.

**Supplementary figure 5:**
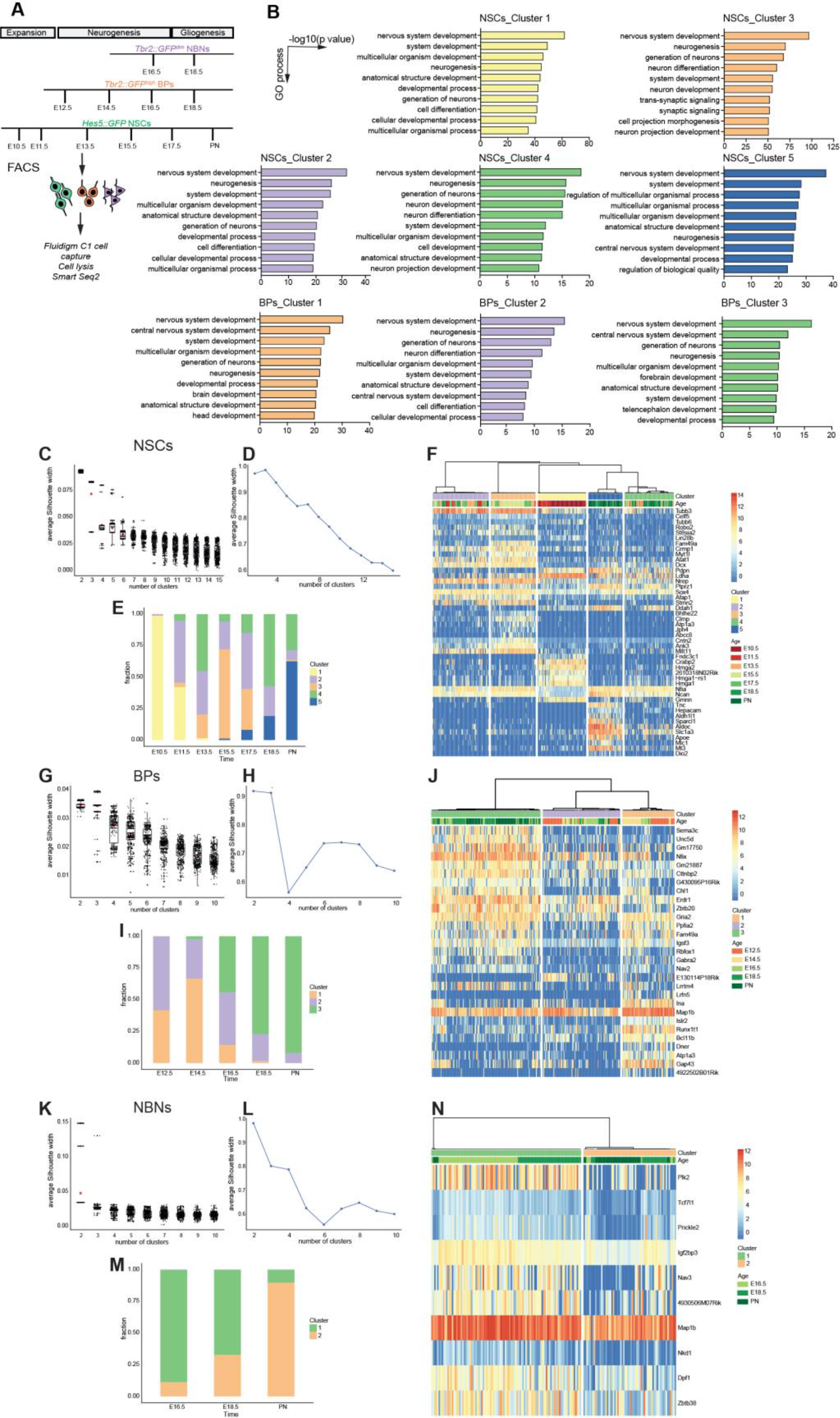
(A) Schematic representation of the experimental approach used for single cell collection used to isolate *Hes5::GFP* and *Tbr2::GFP* cells for single cell sequencing using Fluidigm C1 platform. (B) Gene Ontology (GO) analysis of the biological process in different clusters of NSC’ and BP single cells. Metacore Software was used, -log10(p value) is indicated. The analyses were performed only on the clusters which are composed by 50 or more genes. (C) Silhouette analysis where points represent the average Silhouette width of k-means clusters of NSC single cells for each k for a random initial number. For each k, 500 k-means clustering applied with different initial values. (D) Silhouette coefficient of hierarchal clustering of the assignment matrix of NSC single cells for different k. (E) Bar plot shows the fractions of NSC cells at each cluster at different time points. (F) Signature genes identified for each NSC single cell cluster. (G) Silhouette analysis where points represent the average Silhouette width of k-means clusters of NSC single cells for each k for a random initial number. For each k, 500 k-means clustering applied with different initial values. (H) Silhouette coefficient of hierarchal clustering of the assignment matrix of BP single cells for different k. (I) Bar plot shows the fractions of BP cells at each cluster at different time points. (L) Signature genes identified for each BP single cell cluster. (M) Silhouette analysis where points represent the average Silhouette width of k-means clusters of NSC single cells for each k for a random initial number. For each k, 500 k-means clustering applied with different initial values. (N) Silhouette coefficient of hierarchal clustering of the assignment matrix of BP single cells for different k. (O) Bar plot shows the fractions of NBN cells at each cluster at different time points. (P) Signature genes identified for each NBN single cells cluster.

**Supplementary figure 6:**
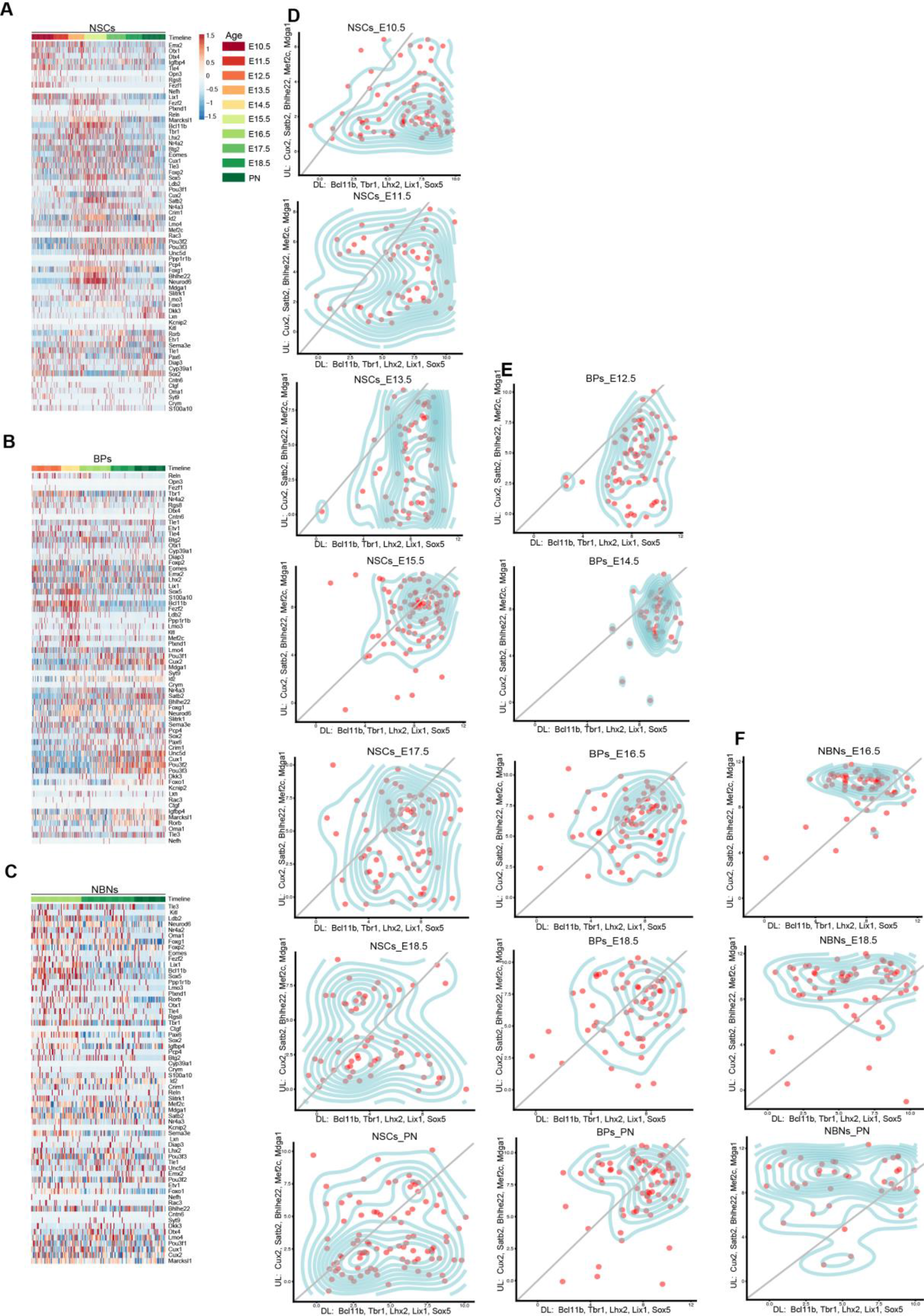
(A) Heatmap of cortical layer markers in NSC single cells, based on z-scored log2(TPM) expression values. (B) Heatmap of cortical layer markers in BP single cells, based on z-scored log2(TPM) expression values. (C) Heatmap of cortical layer markers in NBN single cells, based on z-scored log2(TPM) expression values. (D) Temporal distribution of NSC single cells along the deep or upper layer markers. (E) Temporal distribution of BP single cells along the deep or upper layer markers. (F) Temporal distribution of NBN single cells along the deep or upper layer markers. In D-F, X axis: deep layer markers- Bcl11b, Tbr1, Lhx2, Lix1, Sox5 and Y axis- Cux2, Satb2, Bhlhe22, Mef2c, Mdga1.

**Supplementary figure 7:**
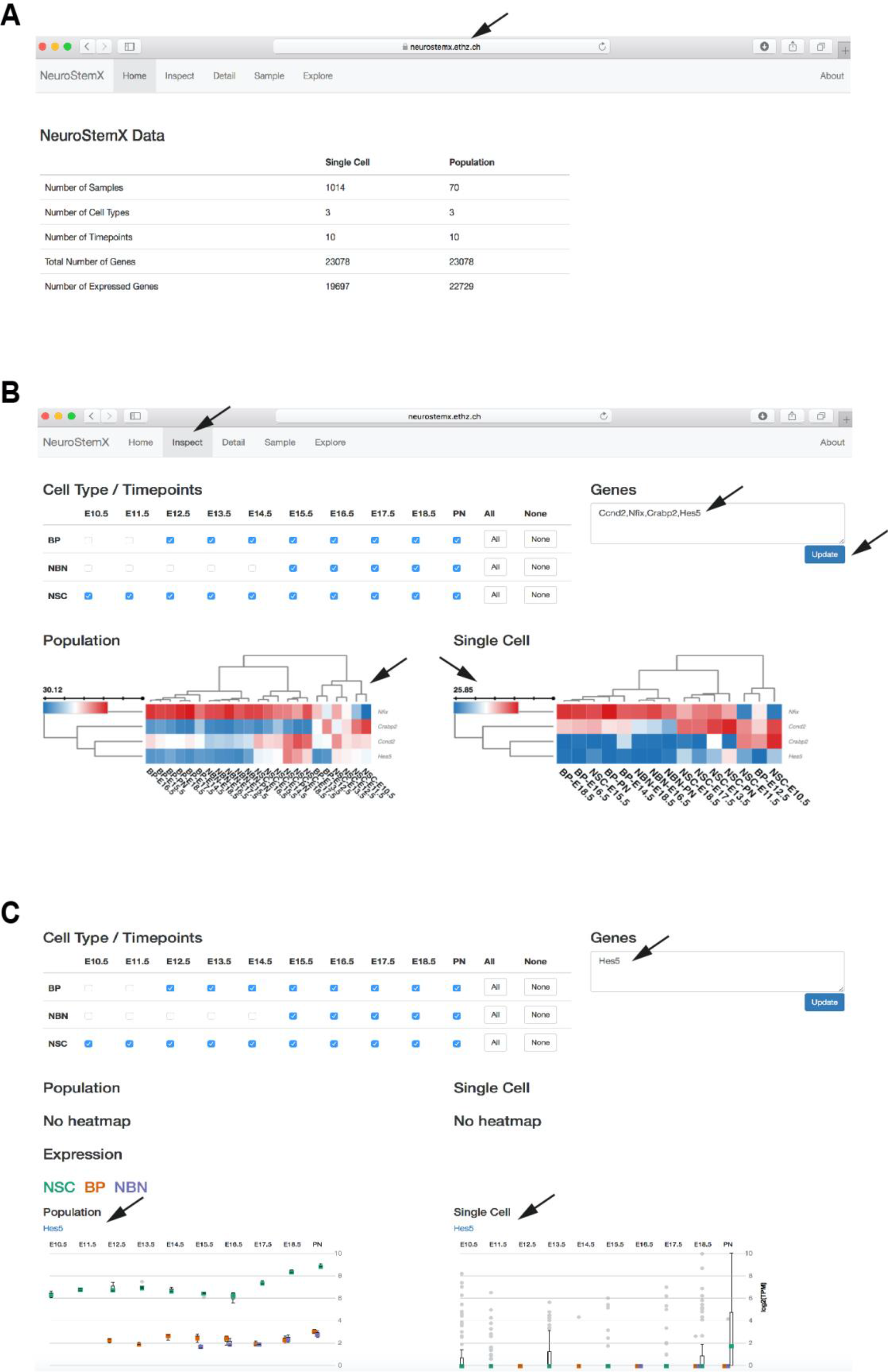
(A) The online browser (http://neurostemx.ethz.ch/) directs to detailed population and single cell RNA sequencing analyses for NSCs, BPs and NBNs. (B) Going through the Inspect tab, one can select the time points or genes one is interested in and click on Update. The website in real-time processes the request and displays the desired heatmaps. (C) Example showing a query for a single gene, here, *Hes5*, yields two types of plots- population and single cell for all cell types. (The data are in log2(TPM), color code as mentioned in the key).

## Table legends

Table 1: This table corresponds to Figures 1 and S1.

Tab1- List of highly variable signature genes from PCA in Figure 1D. Tab2- List of genes in heatmap of Figure 1E.

Tab3- List of DEGs between NSCs and BPs Tab4: List of DEGs between BPs and NBNs Tab5: List of DEGs between NSCs and NBNs

Tab6: Signature genes NSCs versus BPs and NBNs Tab7: Signature genes BPs versus NSCs and NBNs Tab8: Signature genes NBNs versus NSCs and BPs

Table2: This table corresponds to Figures 2, S2 and S3.

Tab1- List of highly variable signature genes from PCA in Figures 2B and S2B, C. Tab2- List of highly variable signature genes from PCA in Figures 2F.

Tab3- List of highly variable signature genes from PCA in Figures 2J and S3C, D. Tab4- List of highly variable signature genes from PCA in Figure 2N and S3C, D.

Tab5: List of genes in NSCs clustered as upregulation, downregulation, transient upregulation, and transient downregulation in Figures S2D-G.

Tab6: List of genes in BPs clustered as downregulation and upregulation in Figure S3E. Tab7: List of genes in NBNs clustered as downregulation and upregulation in Figure S3F.

Table 3: This table corresponds to Figure 3.

Tab1: Motif activity scores for PCA in Figure 3B.

Tab2: List of motifs with fraction of total variance captured by each motif on the first 2 principal components, from Figure 3C.

Tab3: List of predicted motif interaction with interaction likelihood scores in Figure 3D, E. Tab4: Motif activity scores for PCA in Figure 3F.

Tab5: List of motifs with fraction of total variance captured by each motif on the first 2 principal components, from Figure 3G.

Tab6: List of predicted motif interaction with interaction likelihood scores in Figure 3H, I.

Table 4: This table corresponds to Figures 4 and S5.

Tab1-6: Signature genes and summary from 5 NSC clusters identified in Figures 4D and S5F.

Tab7-10: Signature genes and summary from 3 BP clusters identified in Figures 4H and S5J.

Tab11-12: Signature genes and summary from 2 NBN clusters identified in Figures 4L and S5N.

Table 5: This table corresponds to Figures 4 and S5.

Tab1-5: Gene Ontology analyses for 50 enriched categories in 5 NSC clusters identified in Figures 4D and S5F.

Tab6-8: Gene Ontology analyses for 50 enriched categories in 3 BP clusters identified in Figures 4H and S5J.

Table 6: This table corresponds to Figures 6.

Tab1: List of genes considered as markers of stemness, gliogenic and neurogenic phases of NSCs, in Figure 6A-C.

## STAR★METHODS

Detailed methods are provided in the online version of the paper and include the following:

- KEY RESOURCES TABLE
- CONTACT FOR REAGENTS AND RESOURCE SHARING
- EXPERIMENTAL MODELS AND SUBJECT DETAILS

- Mice strain
- METHOD DETAILS

- Tissue preparation and fluorescence-assisted cell sorting
- RNA isolation and RNA-sequencing
- Tissue preparation, immunocytochemistry and immunohistochemistry
- QUANTIFICATION AND STATISTICAL ANALYSIS
- DATA AND SOFTWARE AVAILABILITY

## STAR**★**METHODS

### KEY RESOURCES TABLE

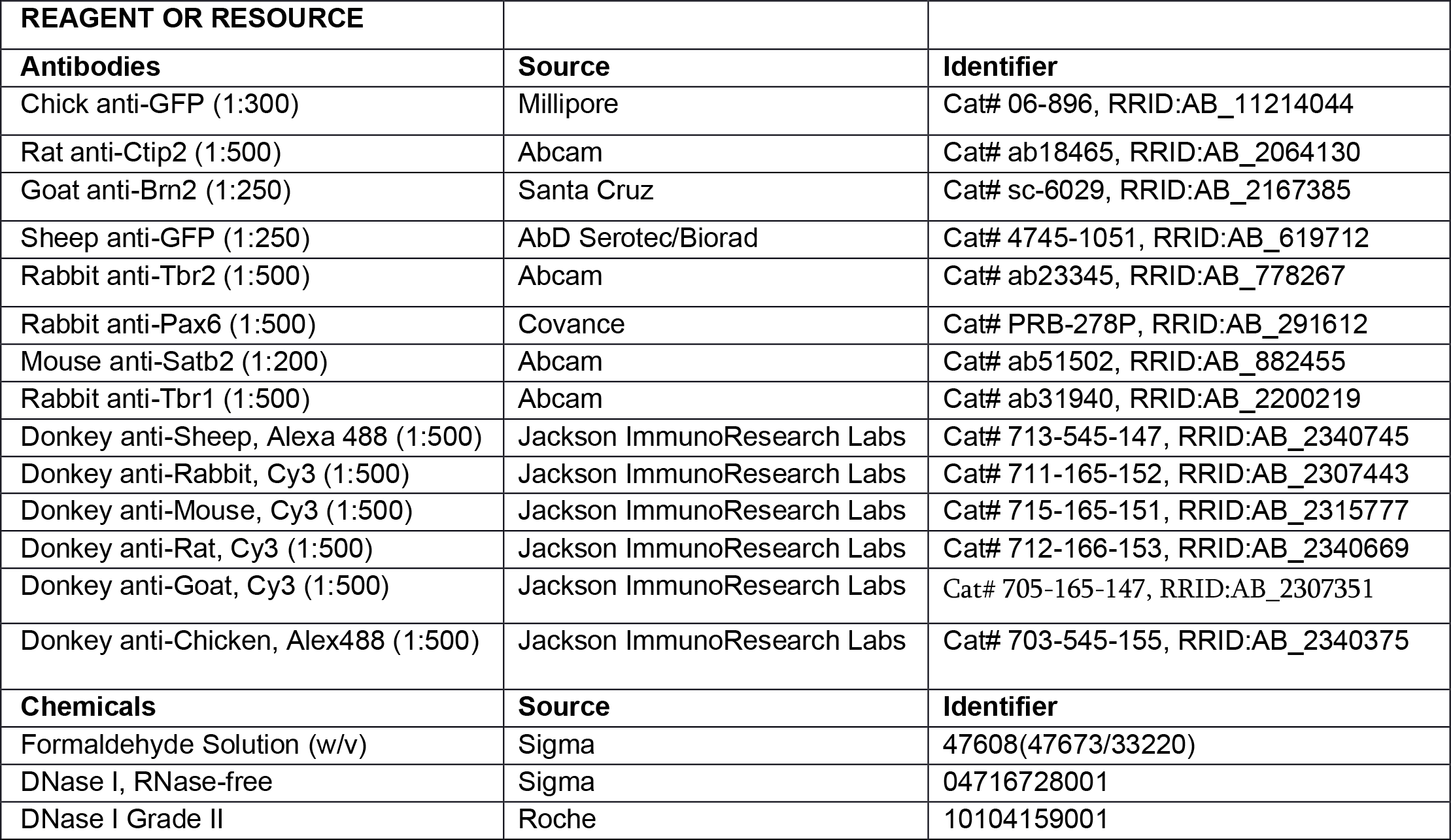
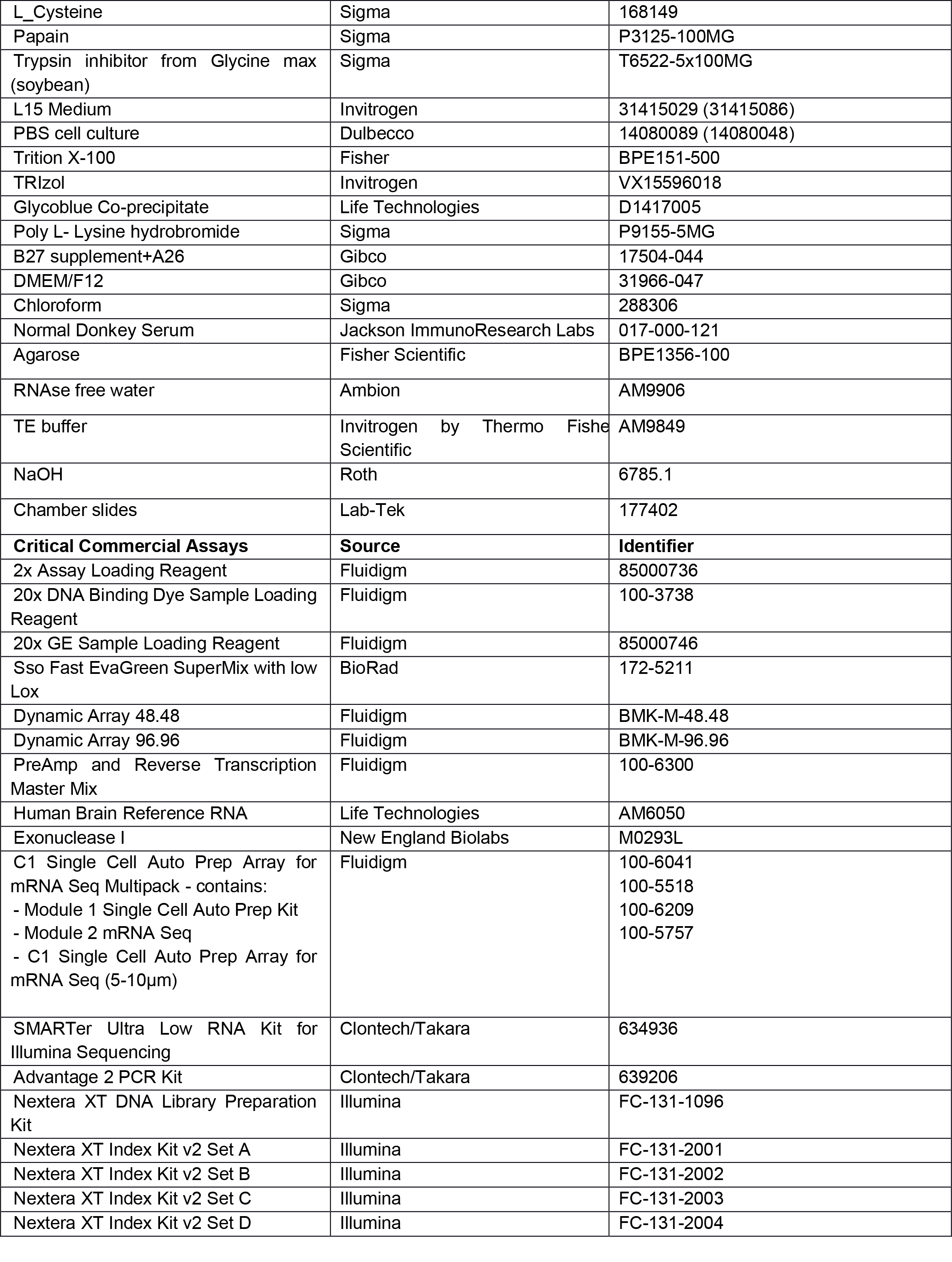
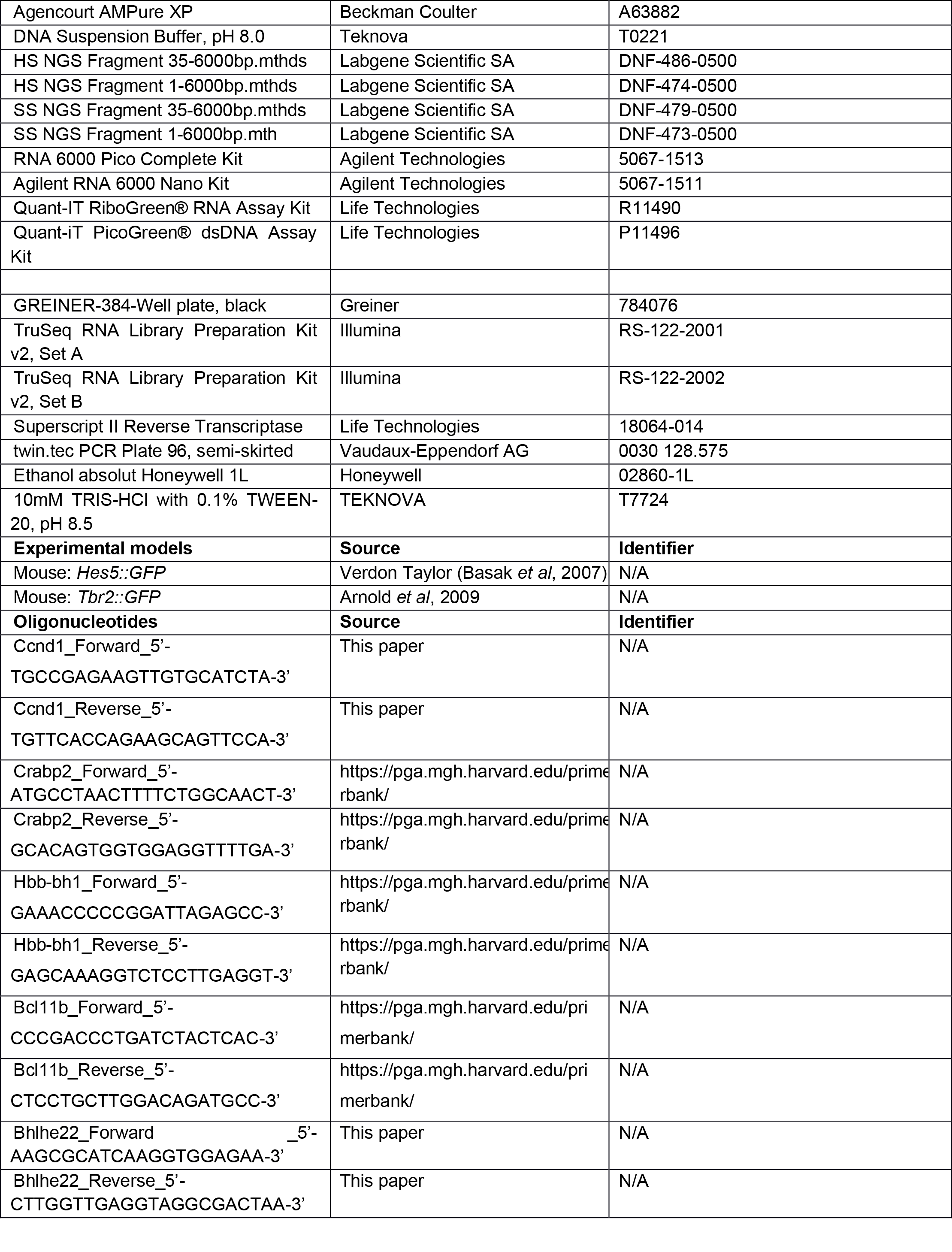
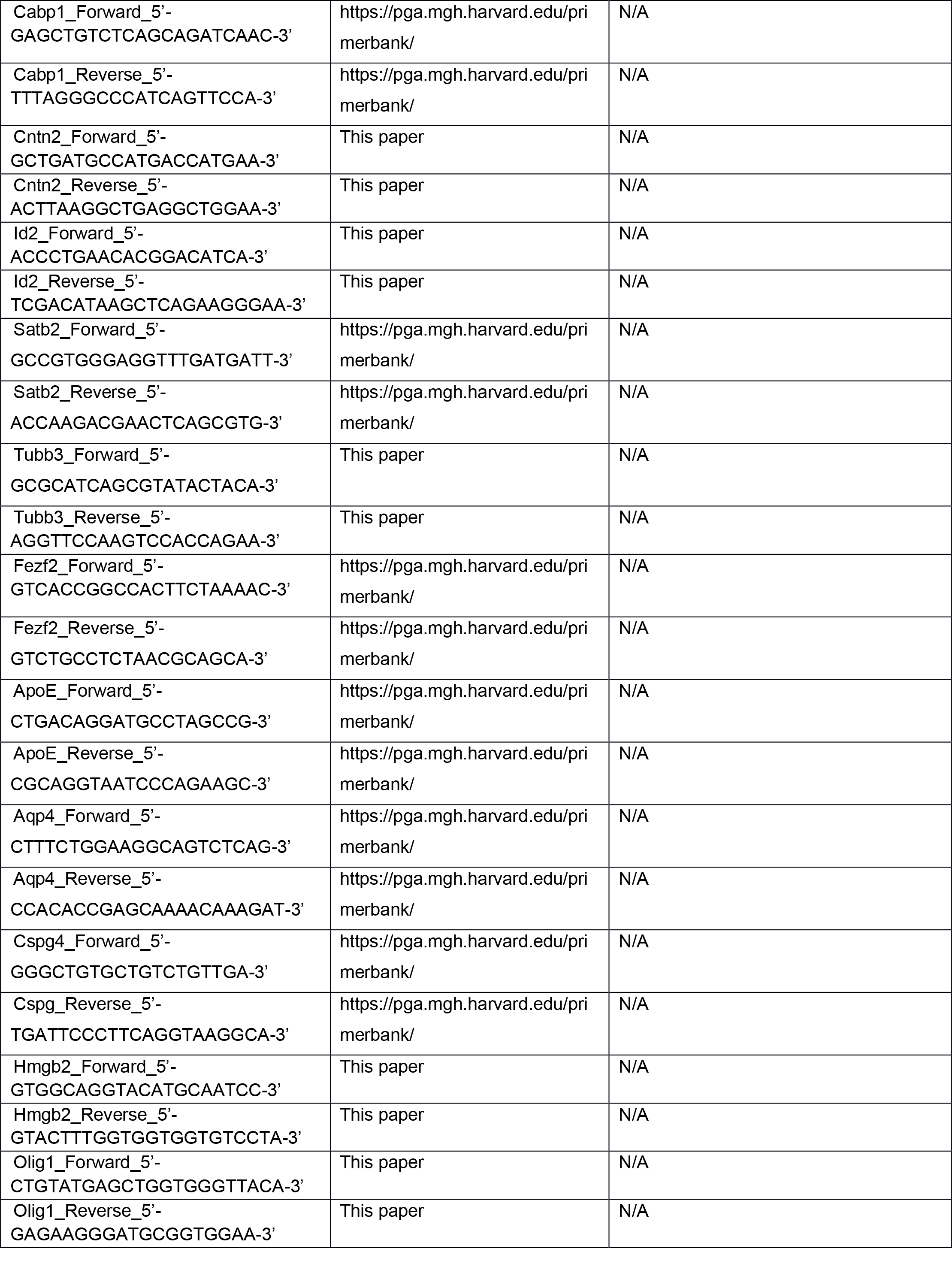
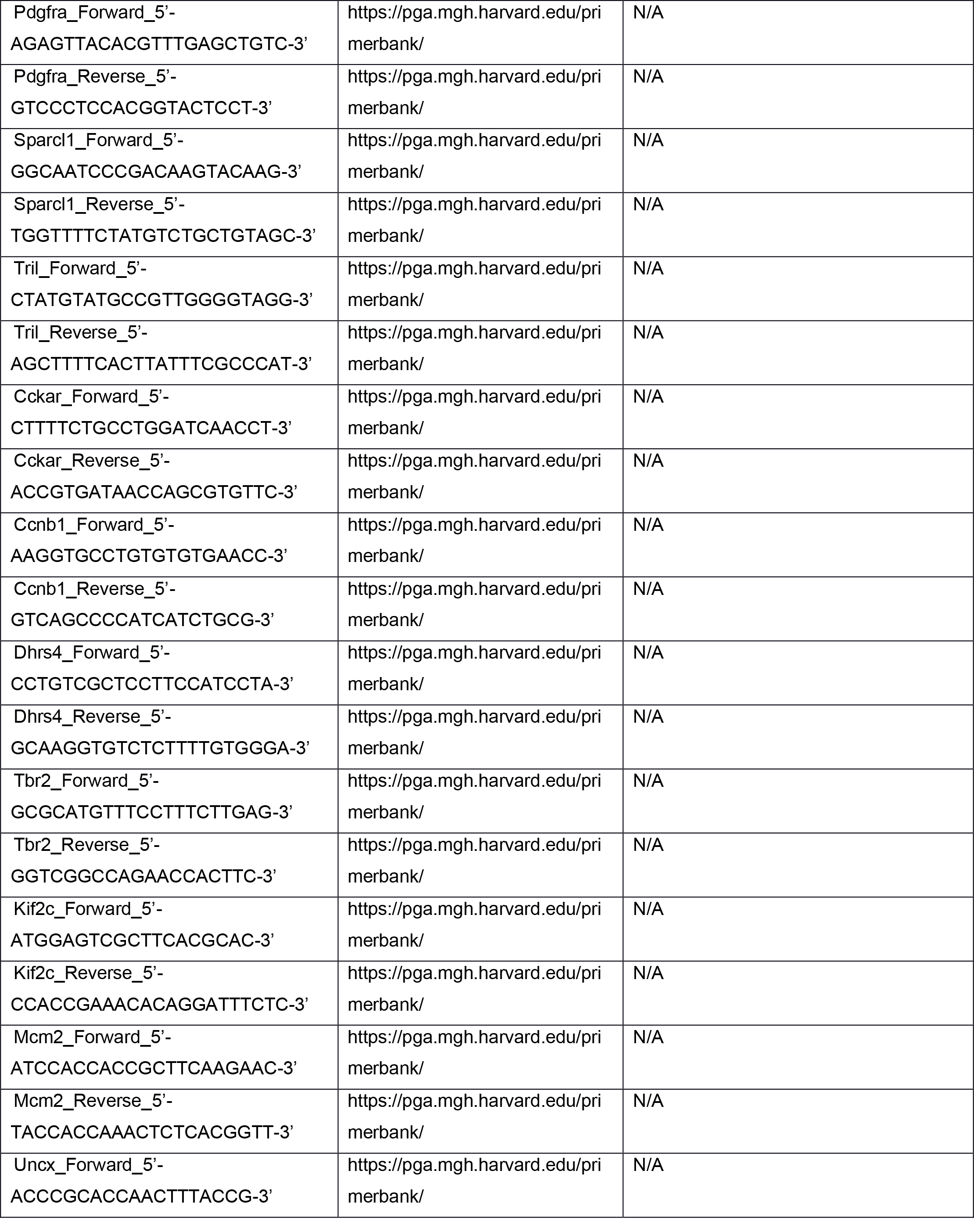
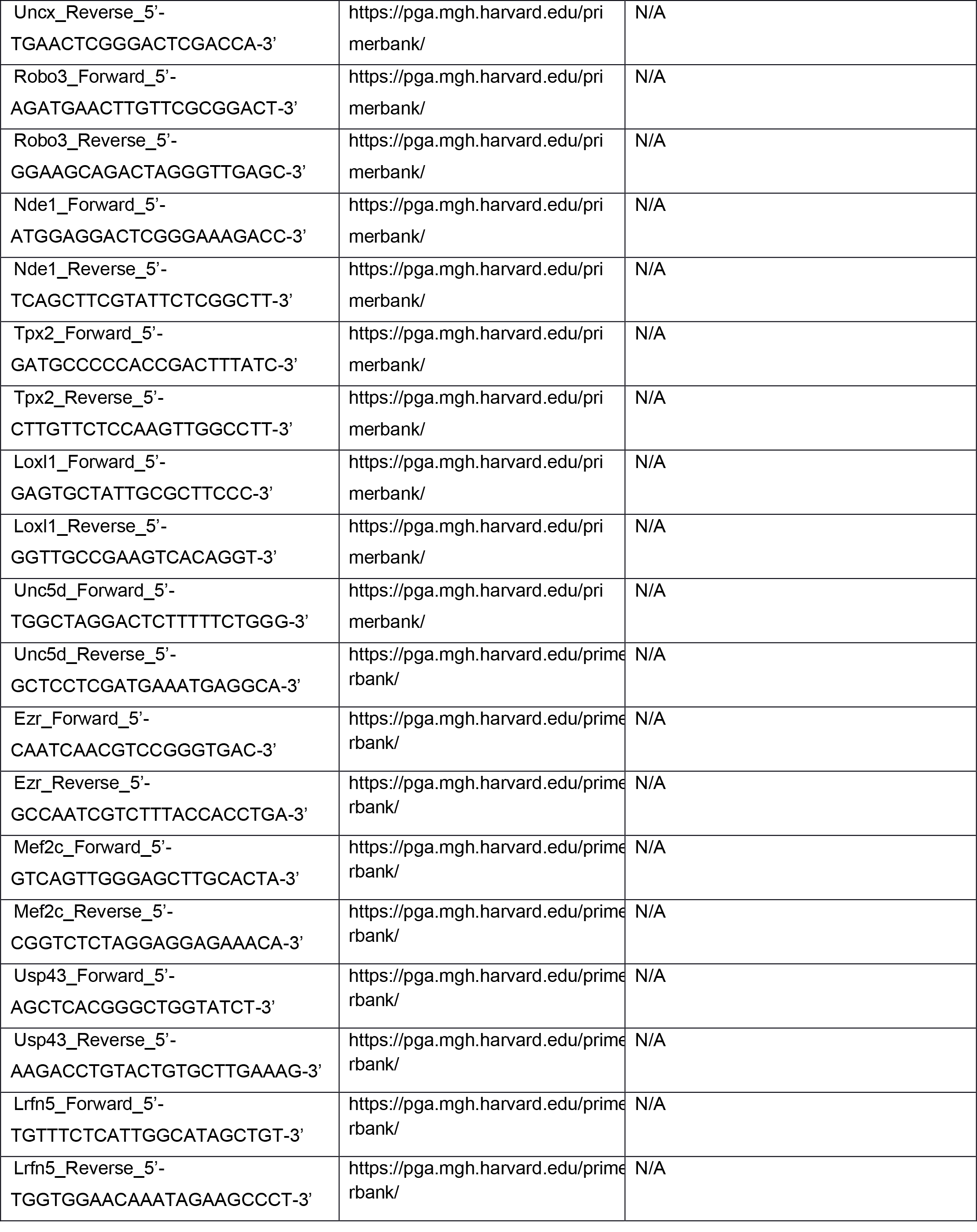
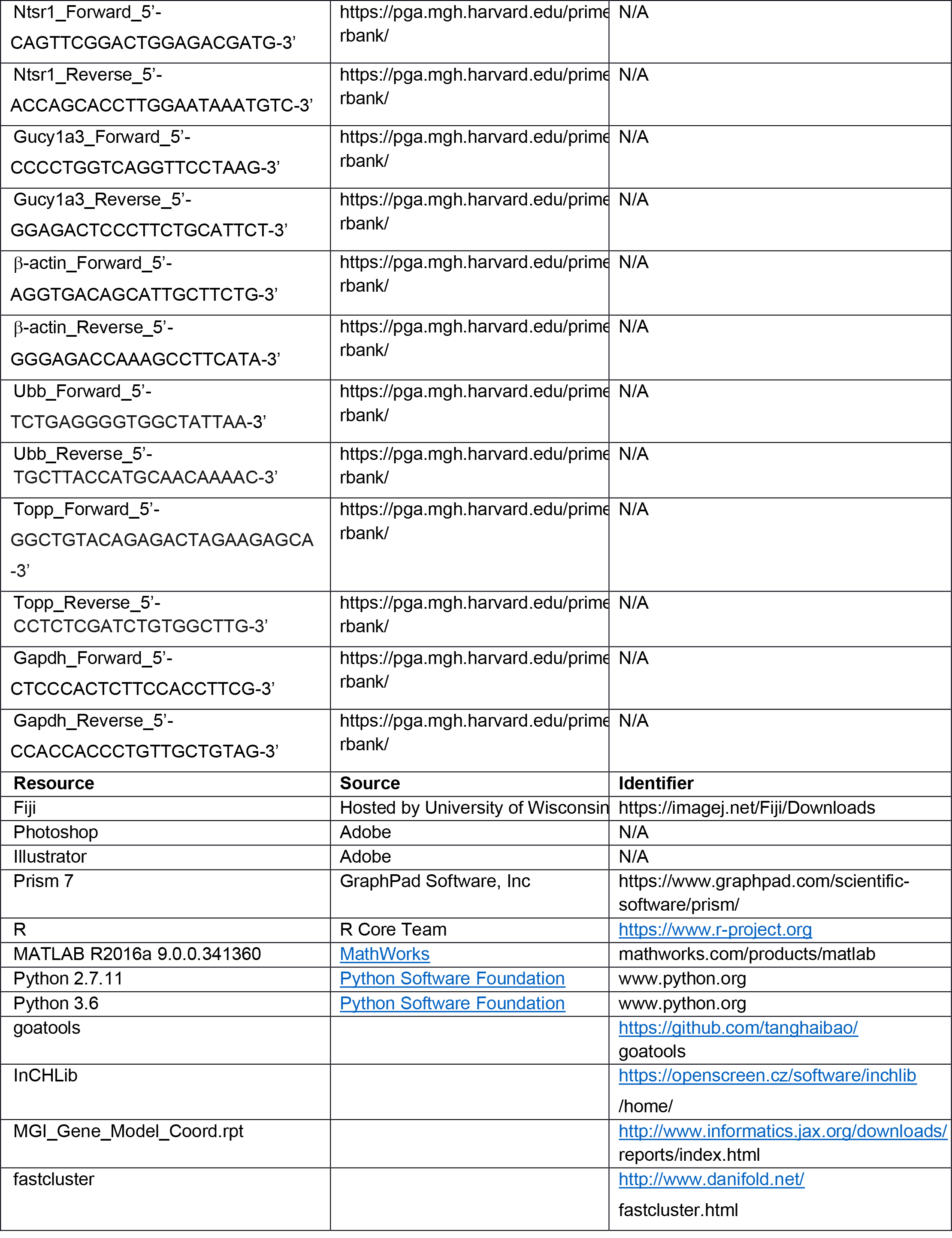
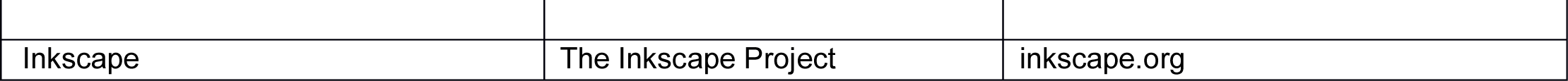

### CONTACT FOR REAGENT AND RESOURCE SHARING

Further information and requests for resources and reagents should be directed to and will be fulfilled by the Lead contact, Verdon Taylor.

### EXPERIMENTAL MODEL AND SUBJECT DETAILS

*Hes::GFP (*Basak *et al,* 2007) and *Tbr2::GFP (* Arnold *et al,* 2009) transgenic lines have been described previously. Mice were maintained on a 12-hr day-night cycle with free access to food and water under specific pathogen-free conditions and according to the Swiss federal regulations. All procedures were approved by the Basel Cantonal Veterinary Office (license number ZH_Tay).

### METHOD DETAILS

#### Tissue preparation and fluorescence assisted cell sorting (FACS)

Dorsal cortices from embryonic day (E10.5) to postnatal day 1 (PN) were micro- dissected and dissociated into single cell suspensions using Papain and Ovo-mucoid mix (as described previously Giachino *et al*, 2009). Cells were washed with L15 medium and FAC-sorted for GFP positive NSCs using FACSariaIII (BD Biosciences) derived from *Hes5::GFP* transgenic embryos for NSCs and *Tbr2::GFP* transgenic embryos for BPs and NBNs. For each time point, 3-4 biological replicates were generated.

#### RNA Isolation and RNA-sequencing

Total RNA was isolated from FAC-sorted GFP positive cells from *Hes5::GFP* and *Tbr2::GFP* transgenic lines using TRIzol reagent. A time course was performed with NSCs, BPs and NBNs isolated at each time point during development from E10.5 to postnatal day 1 (PN), or as specified in the figure 1A. Samples were analyzed for their integrity and concentration using Agilent 2100 Bioanalyzer and Quant-IT RiboGreen RNA Assay Kit. Sequencing libraries were prepared with the Illumina TruSeq RNA Library Prep Kit v2 according to Illumina’s instructions. After quality control (Fragment Analyzer, AATI) libraries were pooled and loaded on an Illumina flow cell for cluster generation (HiSeq SR Cluster Kit v4 cBot). Libraries were sequenced SR50 on the HiSeq 2500 system (HiSeq SBS Kit V4) following the manufacturer’s protocols.

#### Single cell RNA-sequencing

Single cell capture, lysis and cDNA preparation was performed with the Fluidigm C1 system. Cells were loaded on a microfluidic C1 Single Cell Auto Prep Array for mRNA Seq (5-10µm), and capture efficiency evaluated using microscopy. Lysis, reverse transcription, and cDNA amplification was performed with the SMARTer Ultra Low RNA Kit for Illumina Sequencing (Clontech/Takara) according to Fluidigm’s guidelines for single-cell RNA-seq on the C1 system. cDNA was harvested, profiles checked on the Fragment Analyzer (AATI) and their concentration determined using Quant-iT PicoGreen dsDNA Assay Kit. For subsequent library preparation using Nextera XT DNA library preparation kit (Illumina) following the Fluidigm manual, cDNAs were normalized to 0.3 ng/µl. Libraries were pooled and sequenced SR75 on an Illumina NextSeq 500 system (75 cycles High Output v2 kit).

#### qPCR validation

Total RNA was isolated from FAC-sorted GFP positive cells from *Hes5::GFP* (E11.5, E15.5 and E18.5) and *Tbr2::GP (*E13.5 BPs, E15.5 BPs and E15.5 NBNs) transgenic embryos using TRIzol reagent. Independent biological replicates were generated for qPCR validation. Samples were analyzed for their integrity and concentration using Agilent 2100 Bioanalyzer and Quant-IT RiboGreen RNA Assay Kit. DNase treatment was done using Roche DNase kit and cDNA prepared using the PreAmp and Reverse Transcription Master Mix from Fluidigm. Deltagene Assay primers (Fluidigm) and EvaGreen (BioRad) were used for real-time qPCR. Gene expression was assayed using Dynamic Array IFC chips and the BioMark system (Fluidigm). Fluidigm real-time PCR analysis software was used to calculate cycle threshold (Ct) values for each qPCR.

#### Tissue preparation and immunohistochemistry

*Hes5::GFP* and *Tbr2::GFP* positive brains at E17.5 were isolated and fixed with 4% PFA in 0.1M phosphate buffer (PBS). Brains were embedded in 3% agarose, sectioned

40 m thick using a Vibrotome. Sections were mounted in mounting media containing diazabicyclo-octane (DABCO; Sigma) as an anti-fading agent on SuperFrost glass slides and visualized using Zeiss Apotome 2 microscope.

#### Adherent NSC culture *in vitro* and immunocytochemistry

Primary NSCs were isolated from E13.5 dorsal cortices from *Hes5::GFP* transgenic embryos and BPs, NBNs were isolated at E16.5 from *Tbr2::GFP* transgenic embryos. Following FAC-sorting, the cells were seeded in 100 μg/ml Poly L-Lysine pre-coated 8- well Lab-Tek chamber slides and cultured in DMEM/F12 + Glutamax medium (with 2% B27). The cells were incubated for 1h at 37°C, 5% CO2. The cells were fixed with 4% PFA, at RT for 15minutes and blocked with 5% Normal donkey serum and 0.1% Triton X-100. Primary antibody incubations were performed overnight at 4°C. Secondary antibody incubations were performed at RT, for 1h. The cells were incubated with 1:1000 Dapi for 30 minutes at RT and rinse with PBS. Slides were mounted with DABCO and imaged using Zeiss Apotome 2 microscope.

### QUANTIFICATION AND STATISTICAL ANALYSIS

Images taken by Zeiss Apotome 2 were processed with FIJI software. Contrast and image size of IF images were adjusted with Adobe photoshop. Expression profiles of genes of interest were produced in R. Bar graphs were generated by GraphPad Prism 7. All figures were made in Adobe Illustrator CS6.

Sample size is mentioned in the excel sheets for the quantifications. For FACS analysis, for *Hes5::GFP* transgenic embryos, only the bright GFP positive cells were collected. For *Tbr2::GFP* transgenic embryos, both bright and dim GFP positive cells were collected and analyzed. For IF images, three fields of views were analyzed and quantified per sample. Unpaired t-tests were used qPCR validation experiments. The cut-off value for statistical significance were indicated in corresponding figure legend.

### DATA AND SOFTWARE AVAILABILITY

The RNA sequencing datasets have been deposited in Gene Expression Omnibus (GEO) with accession number GEO:

GSE134688 (https://www.ncbi.nlm.nih.gov/geo/query/acc.cgi?acc=GSE134688 and GSE134738 (https://www.ncbi.nlm.nih.gov/geo/query/acc.cgi?acc=GSE134738).

#### Read mapping and data preprocessing

Reads from single cell and cell population mRNA-Seq were mapped to the transcriptome (GENCODE Release M2 GRCm38.p2) with kallisto 0.43.0[*]. The option --pseudobam was used to save the pseudoalignments to transcriptome in BAM file. The reads mapping to multiple transcripts were uniformly distributed. To obtain the expression per transcript, we first divided the number of reads mapping to each transcript by the length of the transcript in nucleotides and then transform the length-normalized read counts in transcript per million (TPM). Gene expression was obtained by summing for each gene the TPM of the transcripts corresponding to the gene. Promoter expression was obtained by summing for each promoter the length-normalized count of the transcripts associated with the promoter and then transformed in TPM. We added a pseudo-count of 0.5 to express transcript, gene, and promoter expression in logarithmic space (log2(TPM+0.5)). For the population mRNA-Seq, we computed replicate averages in log2(TPM+0.5). The method used is adapted from Bray et al, 2016 (Bray et al., 2016a; b).

#### Differentially expressed genes in different cell types

A pairwise comparison between each two cell types is applied using tximport and Deseq2 packages in R. Next, the first 50 top DEGs (differentially expressed genes) for each cell type has been selected considering fold change of more than 2 and adjusted p-value less than 1e-3. Finally, the common DEGs of each cell type is used for visualization. The complete list of DEGs of each comparison is given in (SI. 1 excel sheet).

The goal of our analysis was to find the most optimal marker genes. That is, if we were to only make gene expression measurements of a few genes (using qPCR for example), those that give us the most information about the sample. When we are only interested in knowing whether the sample belongs to one of two classes (e.g., NSC vs non-NSC), this information content is given by the conditional entropy described below. Hence, we use it as a score to find good marker genes. In the derivation we account for the fact that the empirical expression variance from a few data does not necessarily reflect its true variance by using a prior that makes very small and very large variances unlikely.

Assuming that the probability *P*(*x* |*w*, μ) to measure log-expression *x* of a gene follows a Gaussian distribution with mean μ and inverse variance *w*, and using a uniform prior for μ and a gamma-distribution prior 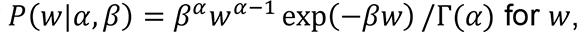, we find the likelihood of getting a set of measurements *D*^*c*^ = (*x*_1_, *x*_2_, … , *x*_*n*_^c^) for samples of class *c* to be

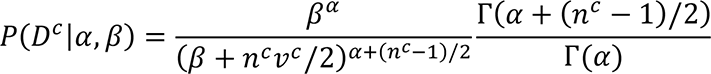

where *v*^*c*^ is the empirical variance of *D*^*c*^. Hence, we numerically find the maximum-likelihood estimates *α*^∗^, *β*^∗^ from maximizing the sum of log-likelihoods across all genes. Finally, the inferred probability distribution of *x* in class *c* is:

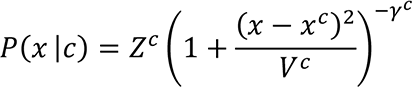

Where 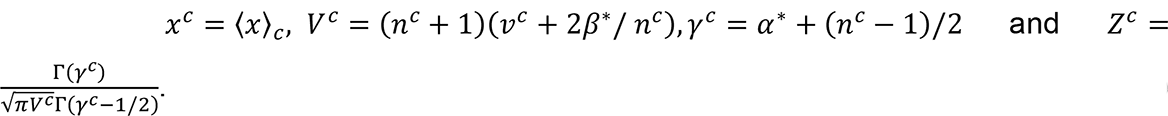. This distribution is approximately Gaussian with variance σ^2^ = *V*^*c*^/(2*γ*^*c*^) which provides us with a more accurate estimate of the true variance of a gene rather than simply taking *v*^*c*^.

Furthermore, we can take the expression of *P*(*x*|*c*) and *P*(*c*) = 1/|*c*| to calculate the conditional entropy *H*(*c*|*x*) = *H*(*x*, *c*) − *H*(*x*). While

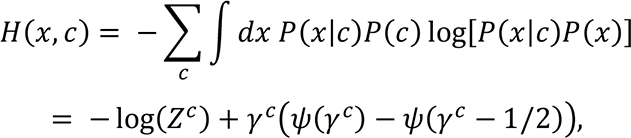

with ψ being the digamma function, has an analytical solution, *H*(*x*) = ∫ *P*(*x*) log(*P*(*x*)) with *P*(*x*) = ∑_*c*_ *P*(*x*|*c*)*P*(*c*) can be calculated through numerical integration.

In an experiment which only measures the expression of a single gene, *H*(*c*|*x*) serves as a measure for how much information the result provides about the class of the sample. With only two classes, we can write *H*(*x*|*c*) = −*p*_*e*_ log *p*_*e*_ − (1 − *p*_*e*_) log(1 − *p*_*e*_), which we can numerically invert to find *p*_*e*_, the probability to falsely classify a sample based on gene expression. The table below summarizes the classes for which we looked for such marker genes:

| | Class 2 | Number of<br>marker genes<br>$p_e < 0.01$ |
| --- | --- | --- |
| **By cell type** |  |  |
| NSC | Non-NSC | 37 |
| BP | Non-BP | 0 |
| NBN | Non-NBN | 136 |
| NSC | BP | 49 |
| NSC | NBN | 469 |
| BP | NBN | 249 |
| **NSC by phase** |  |  |
| Expansion<br>NSC | Non-expansion<br>NSC | 222 |
| Neurogenesis<br>NSC | Non-neurogenesis<br>NSC | 4 |
| Gliogenesis NSC | Non-gliogenesis NSC | 102 |
| Expansion NSC | Neurogenesis NSC | 207 |
| Expansion NSC | Gliogenesis NSC | 759 |
| Neurogenesis<br>NSC | Gliogenesis NSC | 117 |
| **Neurogenic (E12.5-16.5) NSC by day** |  |  |
| E12.5 | Other days | 0 |
| E13.5 | Other days | 3 |
| E14.5 | Other days | 0 |
| E15.5 | Other days | 1 |
| E16.5 | Other days | 50 |
| E12.5-E13.5 | E14.5-E16.5 | 9 |
| E12.5-E14.5 | E15.5-E16.5 | 16 |
| **BP by day** |  |  |
| E12.5 | Other days | 248 |
| E13.5 | Other days | 0 |
| E14.5 | Other days | 29 |
| E15.5 | Other days | 0 |
| E16.5 | Other days | 1 |
| E17.5 | Other days | 3 |
| E18.5 | Other days | 1 |
| PN | Other days | 38 |
| E12.5-E13.5 | E14.5-PN | 54 |
| E12.5-E14.5 | E15.5-PN | 83 |
| E12.5-E15.5 | E16.5-PN | 0 |
| E12.5-E16.5 | E17.5-PN | 15 |
| E12.5-E17.5 | E18.5-PN | 4 |
| **Other** |  |  |
| BP<br>E12.5-E14.5 | NBN | 354 |
| BP E15.5-PN | NBN | 650 |

#### Selection of highly variable genes

To select the most highly variable genes, we have defined a score for each gene based on the contribution of each gene on each principal component and the variance that each component explains considering the first two components, as following:

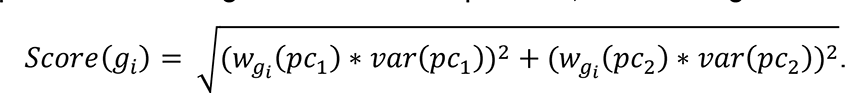

Where *w*_*gi*_ (*pc*_*k*_) refers to the weight (contribution) of gene *pc*_*k*_ and *varpc*_*k*_ denotes the percentage of variance that is covered by *pc*_*k*_. Next, the first 2000 genes with the highest scores are selected as the highly variable genes, HVGs.

#### Clustering of single cells

First, single cells at each time point are clustered by applying 500 times k-means clustering to avoid the dependency of k-means clustering on the random initialization number (seed value). To implement k-means clustering, *clustering* package considering Euclidean distance as metric in R is used. Next, the assignment matrix is estimated based on the frequency of observing each two single cells in the same clustering at each iteration. Next, the hierarchal clustering is used to sort the assignment matrix using Euclidean distance as metric and *ward. D2* as method in *hclust* function in R.

#### Selection of differentially expressed gene in single cells

Kruskal-Wallis non-parametric test is used to select differentially expressed genes and genes with adjusted p-value of less than 1e-3 are considered as significantly differentially expressed genes.

#### Visualization

PCA is applied using *prcomp* function in R after centering the log transferred data.

Heatmap are illustrated using *pheatmap* package in R on log transferred data.

#### NeuroStemX Data Exploration Web App

The NeuroStemX data exploration web app makes it possible to navigate data produced in the NeuroStemX project. The site supports viewing data by focusing on one of several parameters: gene list, biological sample, or measurement type (single cell vs. population).

The website allows entry of a list of mouse genes (either as gene symbol or Ensembl ID) to focus on the data acquired for those genes. It alternatively allows viewing data on individual samples. When looking at a sample, a list of genes that have been determined to be outliers are shown. A gene is considered an outlier for a sample if the expression value of the gene either exceeds the 75th percentile + 1.5*iqr or is less than the 25th percentile - 1.5*iqr, where percentiles and inter-quartile range are computed based on the expression values for the given gene over all samples within the measurement type (single cell or population).

When viewing all data for a measurement type, data is displayed using hierarchical clustering. The InCHLib widget displays the clustered data. Clustering is performed using the fastcluster package in python with distance (both row and column) calculated using the Euclidean metric and linkage (both row and column) performed using Ward’s method (Mullner, 2013; Skuta et al., 2014).

The site supports performing gene ontology enrichment analysis either locally, using goatools, or with PANTHER. For local enrichment analysis, we use the MGI_Gene_Model_Coord annotations based on the GRCm38 assembly (Mi et al., 2017).

**Goatools**: https://github.com/tanghaibao/goatools

Haibao Tang *et al*. GOATOOLS: Tools for Gene Ontology. Zenodo. 10.5281/zenodo.31628.

#### MGI_Gene_Model_Coord.rpt

http://www.informatics.jax.org/downloads/reports/index.html

